# Locus-specific gene-context interactions improve polygenic prediction

**DOI:** 10.64898/2026.08.26.746823

**Authors:** Renée Fonseca, Christa Caggiano, Manuela Costantino, Ophelia Dominguez, Eimear E. Kenny, Andy Dahl

## Abstract

Polygenic scores (PGS) are a primary output of large-scale genetic studies and are being deployed in clinical and non-clinical settings. However, current PGS assume simple additive models that ignore context-specific genetic effects, which likely reduce their accuracy and robustness. To address this, we developed PGSC, a PGS framework to incorporate locus-specific gene-context interaction effects (GxC). Simulations show PGSC is robust under the additive model and outperforms PGS in realistic settings. Using sex, age, and statin treatment status as contexts in UK Biobank, we find that PGSC outperforms PGS on average across 48 traits, with substantial improvement in some cases, such as GxSex for testosterone, GxAge for bilirubin, and GxStatins for LDL cholesterol. PGSC consistently outperforms a simple genome-wide GxC model, ampPGS, which only outperforms PGS when a context uniformly amplifies all genome-wide additive effects. Critically, PGSC improvements replicate across ancestries in the UK Biobank and in an external cohort, the Mount Sinai Million Health Discovery Program. Finally, we test robustness to log-scale phenotypes and find that ampPGS gains vanish, while the locus-specific GxC components in PGSC persist. Overall, PGSC is a simple, robust framework that demonstrates GxC effects can improve out-of-sample PGS prediction and is a step toward precision treatment.

## Introduction

Genome-wide association studies (GWAS) have identified thousands of genetic variants associated with complex traits. Polygenic scores (PGS) aggregate these associations across the genome to predict individual-level phenotypes. A major goal of PGS is to improve clinical care, for example, by identifying high-risk individuals for preventive measures^1^. PGS also hold promise for predicting an individual’s treatment response^2,3^, and recent work has shown that genetic effects can substantially modify responses to common drugs such as statins^4^. However, current PGS remain far less accurate than the theoretical optimum, and they often transfer poorly to new settings, motivating more sophisticated PGS modeling approaches.

One limitation of standard PGS is their assumption of a simple additive model, where all genetic effects act equally in all contexts. Yet non-additive effects due to gene-context interaction (GxC) often explain substantial heritability in complex human traits^4–11^. GxC effects may also partly explain why PGS transfer poorly across contexts such as age^12^, sex^10^, ancestry^13,14^, or phenotyping strategy^15^. Together, high GxC heritability and low PGS transferability suggest that incorporating GxC effects should improve PGS robustness and accuracy.

However, less attention has been given to realizing the potential of GxC-aware PGS. Currently, the most common approach is a simple PGS amplification model (ampPGS)^11^, which assumes that context uniformly amplifies or buffers every genetic effect. ampPGS is practical because it requires only one GxC parameter and standard GWAS results, but it cannot capture locus-specific GxC^5,7,9,16^. More fundamentally, GxC signals are challenging to model, as they often vanish after phenotype transformation^17,18^ or require accounting for heteroskedastic noise^4,5,8^, and they are rarely replicated across cohorts or ancestries^11,19^. Overall, GxC has not yet been shown to improve PGS prediction (Table S1).

Here, we address this gap by developing PGSC, a simple framework to incorporate locus-specific GxC effects into polygenic scores. Using sex, age, and statin treatment status as contexts in the UK Biobank (UKB)^20^, we find that PGSC robustly improves prediction: PGSC outperforms PGS on average across traits and substantially outperforms PGS when GxC heritability is high. PGSC consistently outperforms ampPGS; further, ampPGS gains vanish on the log scale, while PGSC gains largely persist. Importantly, we demonstrate that PGSC gains are robustly transferable by replicating across ancestries within UKB and in the Mount Sinai Million Health Discovery Program (MSM)^21^.

## Results

### PGSC overview

Standard PGS build an additive genetic score by multiplying additive GWAS effects, 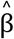, by the genome-wide genotype vector, *g*. PGSC similarly builds a GxC-based PGS using the locus-specific GxC GWAS effects, 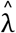, and then adds this to the standard PGS:

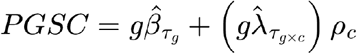

where the first term is the standard PGS with additive p-value threshold *τ*_*g*_ and the second term is a GxC score with GxC p-value threshold *τ*_*g*×*c*_. ρ_*c*_ are the key GxC tuning parameters and depend on the context *c*; for simplicity, we assume the context is binary with values *a* and *b*, so *c* ∈ {*a, b*}. Importantly, learning ρ_*c*_ = 0 enables PGSC to simplify to the PGS under the additive homoskedastic model. We use clumping+thresholding (C+T) to learn *τ*_*g*_ and *τ* _*g*×*c*_, but any PGS method could be used to tune 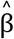 and 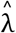 (Methods). PGSC first learns 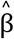 and 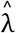 in a GWAS training cohort, then learns *τ*_*g*_, *τ*_*g*×*c*_, and ρ_*c*_ in a tuning cohort, and finally evaluates prediction accuracy in a validation cohort (Fig. 1). PGSC is computationally efficient and practical because it leverages PLINK 2.0^22^ for implementation (Methods).

**Fig. 1.**
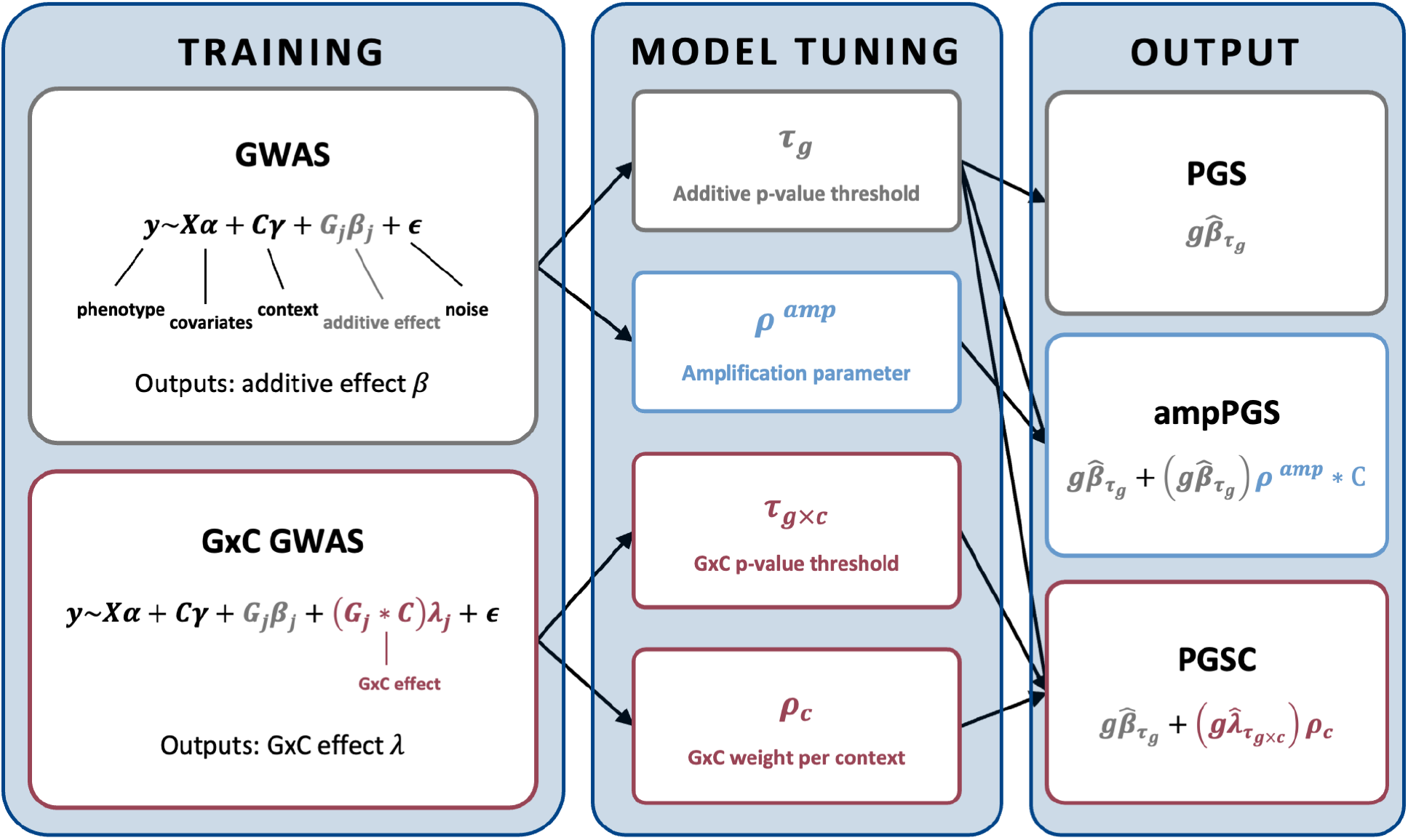
Overview of the PGSC framework. PGSC extends the standard PGS framework by incorporating locus-specific GxC GWAS effects along with additive GWAS effects, scaling the GxC component by an individual’s context. Additive and GxC effect sizes are estimated in a GWAS training cohort; SNP clump + threshold parameters (*τ*_*g*_, *τ*_*g*×*c*_) and context-specific tuning parameters (ρ_*c*_) are optimized in a held-out tuning cohort; predictive accuracy is evaluated in a separate held-out validation cohort. We compare PGSC to the standard PGS and ampPGS, a simpler GxC prediction model based on genome-wide amplification effects.

We might expect ρ_*c*_ = 0 under the additive model, as no GxC is present. However, when unmodeled context-specific noise levels exist (i.e., heteroskedasticity), performance improves by weighting estimates toward the less-noisy context. Concretely, the optimal value for ρ_*c*_ becomes (Note S1):

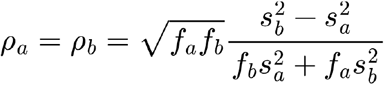

where 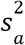 and 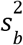 are the residual phenotypic variances in contexts *a* and *b*, respectively, and *f*_*a*_ and *f*_*a*_ = 1 − *f*_*a*_ are the fractions of individuals in each context. This extends the comparison of pooled and context-specific GWAS estimates from Weine et al. 2025^23^ to an optimal weighted average.

We compare PGSC to a simpler model we call ampPGS, which learns a single context-specific weight, ρ^*amp*^, on the standard PGS (Methods):

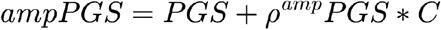

ampPGS cannot model locus-specific GxC because context interacts identically with all genetic effects in the PGS^5,7,9,10,16,24^. This simplifying assumption makes ampPGS convenient, as it does not require performing GxC GWAS, leading to its widespread adoption as an in-sample statistical test for polygenic GxC^11^. Further, PGS accuracy has been shown to differ across contexts such as drug use status^3,4,25^, sex^10,12^, age^12^, socioeconomic status^26^, and phenotype ascertainment strategy^15^. These results suggest that ampPGS should be able to improve out-of-sample PGS prediction.

### PGSC performance in simulations

We compared PGSC to PGS and ampPGS by simulating from the standard polygenic model with additive heritability 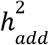 and GxC heritability 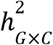 (the proportions of phenotypic variance explained by additive and GxC effects, Methods, Note S2). Under the additive homoskedastic model, where GxC is absent 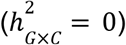, PGSC and PGS performed similarly (Fig. S1). This is crucial in practice because we expect 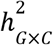 to be near zero for many gene-context pairs. As expected, PGSC performance progressively improves with GxC signal strength, 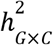 (Fig. 2a).

**Fig. 2.**
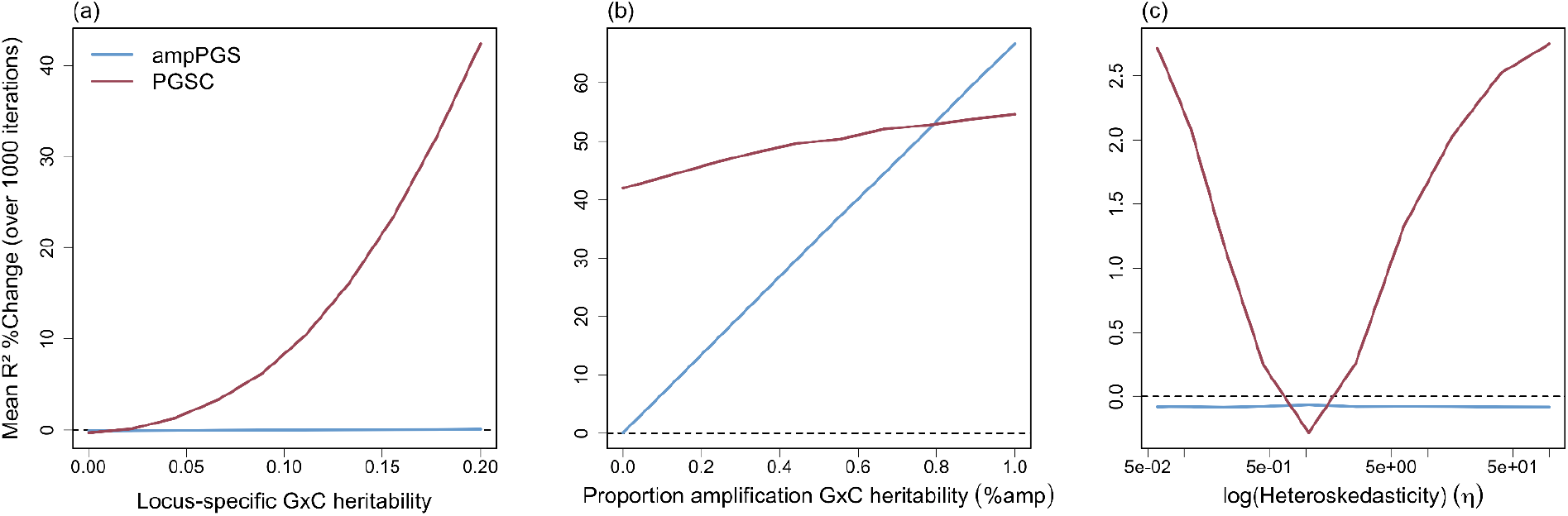
PGSC and ampPGS performance in simulations. (a) GxC effects were simulated as locus-specific (purely random). (b) The proportion of total GxC heritability attributable to amplification GxC effects (%amp) was varied from 0 to 1, with total locus-specific and amplification GxC heritability held constant at 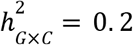. (c) Phenotypes were simulated with no 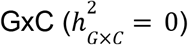 while varying the degree of heteroskedasticity (η). Unless otherwise noted, all simulations used the following baseline parameters: additive heritability 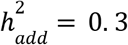, *S* = 10^4^ total SNPs with *S*_*causal*_ = 0. 1 × *S* causal SNPs, p-value thresholds *τ* ∈ {10^−10^, 10^−9^, …, 10^0^} total sample size *N* = 10, 000, and cross-context variance ratio η = 1 (no heteroskedasticity). We ran 1,000 simulation replicates for each parameter setting. We measured prediction accuracy using R^2^ %Change relative to PGS for PGSC (red) and ampPGS (blue).

Our simulations above assumed a form of locus-specific GxC where GxC effects are randomly distributed across loci^27^. Under this model, ampPGS does not outperform PGS even when 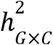 is large, since it cannot capture this locus-specific GxC (Fig. 2a). Next, we introduced genome-wide amplification GxC effects 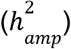, varying the proportion compared to locus-specific GxC (%amp) while holding the total 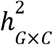 constant (Fig. 2b). PGSC performance grew only slightly with %amp, as expected since its model is agnostic to the form of GxC. Conversely, ampPGS directly scales with %amp; in fact, as %amp approached 100%, its more parsimonious model enabled it to outperform PGSC.

We then simulated an additive model with context-dependent noise levels (i.e., heteroskedasticity). As expected, PGSC partly captured this signal by re-weighting toward the less-noisy context’s additive estimate, thus improving performance over PGS. Conversely, ampPGS could not capture this heteroskedasticity signal (Fig. 2c). Finally, we also found that PGSC improves over PGS as sample size increases or polygenicity decreases, and we confirmed that PGSC is robust to non-uniform context distributions (Fig. S1).

Overall, PGSC outperforms PGS given sufficient GxC signal and/or heteroskedasticity, while ampPGS only succeeds under a particular form of genome-wide GxC.

### PGSC improves prediction in UKB

We evaluated PGSC prediction accuracy in UKB across 48 quantitative traits and three contexts: sex, age, and statin treatment status (Table S2). We built PGSC in 335,461 unrelated White British (WB) individuals (90% for GWAS, 10% for PGS tuning) and evaluated accuracy in 24,951 unrelated European (Euro) individuals (the two largest UKB populations as defined in Dahl et al. 2023^28^, Methods, Table S3). We measure prediction based on the relative change in incremental Pearson R^2^ (R^2^ %Change).

We first examined sex as a context (GxSex), given its well-studied role in statistical genetic interactions^10,29–31^. On average across traits, PGSC improved prediction R^2^ over standard PGS by 5.2% (p = 5.6×10^−69^; Fig. 3a; Table S4). R^2^ %Change varied substantially by trait and was significantly above zero for 24/48 traits (p < 0.05), with the largest gains for sexually dimorphic phenotypes such as testosterone (104.5%) and BMI-adjusted waist-to-hip ratio (WHRadjBMI, 32.3%).

**Fig. 3.**
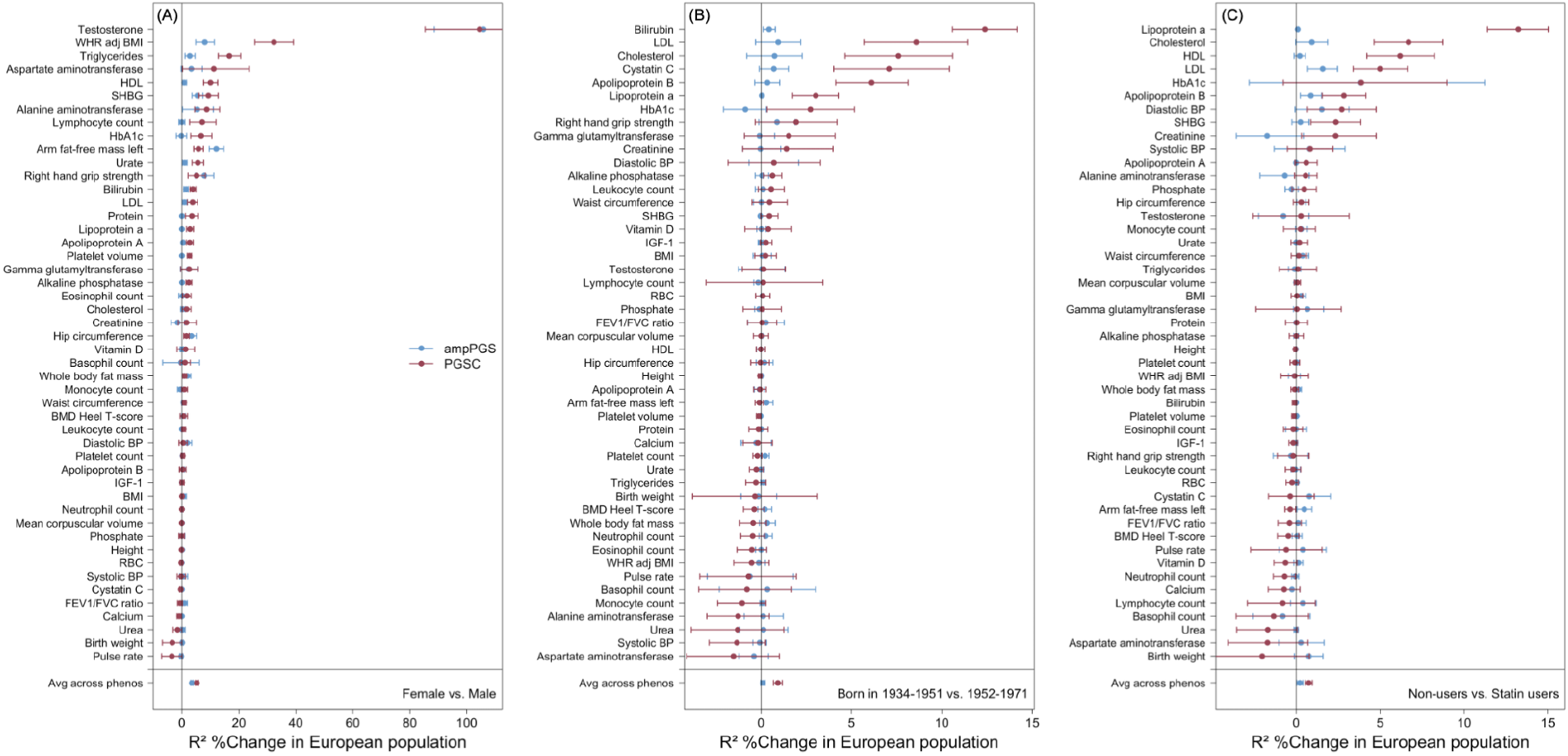
PGSC improves phenotype prediction in UKB. R^2^ %Change for phenotype prediction achieved by PGSC (red) and ampPGS (blue) relative to PGS across 48 quantitative traits, for contexts (A) sex, (B) age, and (C) statin treatment status. The x-axis shows the relative change in incremental R^2^, while the y-axis lists individual phenotypes, with the average improvement across traits shown at the bottom. Error bars represent 95% confidence intervals from 10,000 bootstrap samples. We trained scores on a set of unrelated WB individuals and evaluated them on an independent set of unrelated Euro individuals from UKB.

In comparison, ampPGS improved R^2^ by 3.5% over PGS on average and was individually significant for 16/48 traits (p = 3.9×10^−50^). PGSC outperformed ampPGS on average across traits (R^2^ %Change = 1.7, p = 2.6×10^−12^, Fig. S2), which was individually significant for 15/48 traits at p < 0.05; conversely, ampPGS outperformed PGSC for 6/48 traits at p < 0.05.

We next evaluated age as a context (GxAge). Again, PGSC outperformed PGS on average across traits (R^2^ %Change = 0.93, p = 5.5×10^−14^; Fig. 3b; Table S5), with the largest gains for age-associated biomarkers including bilirubin, LDL, total cholesterol, and cystatin C (R^2^ %Change = 7.1–12.4). In striking contrast to GxSex, ampPGS with GxAge only outperformed the PGS for 2/48 traits at p < 0.05 and did not significantly outperform PGS on average (R^2^ %Change = 0.1, p = 0.11).

Finally, we analyzed statin treatment status as a context (GxStatins)^4^. Statins are a class of LDL cholesterol-lowering drugs commonly prescribed to reduce heart disease^32,33^. PGSC using GxStatins outperformed PGS on average across traits (R^2^ %Change = 0.75, p = 1.9×10^−11^; Fig. 3c; Table S6), with the largest gains for cholesterol-related traits (Lp(a), HDL, LDL; R^2^ %Change = 5.0–13.2). ampPGS with GxStatins significantly outperformed PGS on average, although to a much lesser degree than with GxSex (R^2^ %Change = 0.2, p = 0.02).

We then asked how R^2^ %Change per phenotype related to established parameters of genetic architecture. First, we compared to GENIE estimates of 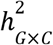, the heritability explained by GxC, and η, the heteroskedastic variance, as reported by Pazokitoroudi et al. 2024 (Table S7)^8^. We found that R^2^ %Change was positively correlated with 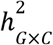 for both PGSC and ampPGS across all three contexts (all 6 have p < 0.05, Ext Data Fig. 1); likewise, R^2^ %Change was positively correlated with the main effect of the context (5/6 have p < 0.05, Fig. S10). We found that PGSC R^2^ %Change increased with η for GxAge (p = 0.02), while ampPGS did not, consistent with the expectation that only PGSC can capture heteroskedasticity. The PGSC context weights behaved as expected: the difference (ρ_*a*_ − ρ_*b*_) was correlated with 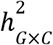 (p < 0.05 for all three contexts), and the sum (ρ_*a*_ + ρ_*b*_) was correlated with η for GxAge (p = 0.05, Ext Data Fig. 2). We also considered a measure of locus-specific GxC, the number of GxC GWAS loci, which correlated with PGSC R^2^ %Change for GxSex and GxAge (p < 0.05, Fig. S3) and to a lesser degree with ampPGS R^2^ %Change for GxSex. Overall, these results show how PGSC can capture amplification, locus-specific GxC, and/or heteroskedasticity, which vary widely across traits and contexts depending on genetic architecture.

### PGSC improves cross-population prediction in UKB

PGS are well-known to perform worse for individuals with ancestries more distant from the GWAS population^13,14^. GxC is one of several potential explanations for this gap. We tested this by asking if PGSC trained in the WB population could improve prediction in 7,486 African (Afr) and 2,459 East Asian (Asn) unrelated individuals (the third and fourth largest populations defined in Dahl et al. 2023^28^) relative to PGS. On average across traits, PGSC consistently outperformed PGS across populations and contexts (avg R^2^ %Change = 5.40, all p < 0.001; Fig. 4, Figs. S4-5; Tables S8-13). ampPGS improved average performance in the Afr and Asn populations for GxSex to a lesser degree than PGSC, but ampPGS did not show improvements on average for GxAge or GxStatins (Fig. 4; Figs. S4-5). Per-trait estimates of R^2^ %Change were significantly correlated with estimates in Euro for PGSC and ampPGS (Table S14), though they were noisier due to the smaller sample sizes.

**Fig. 4.**
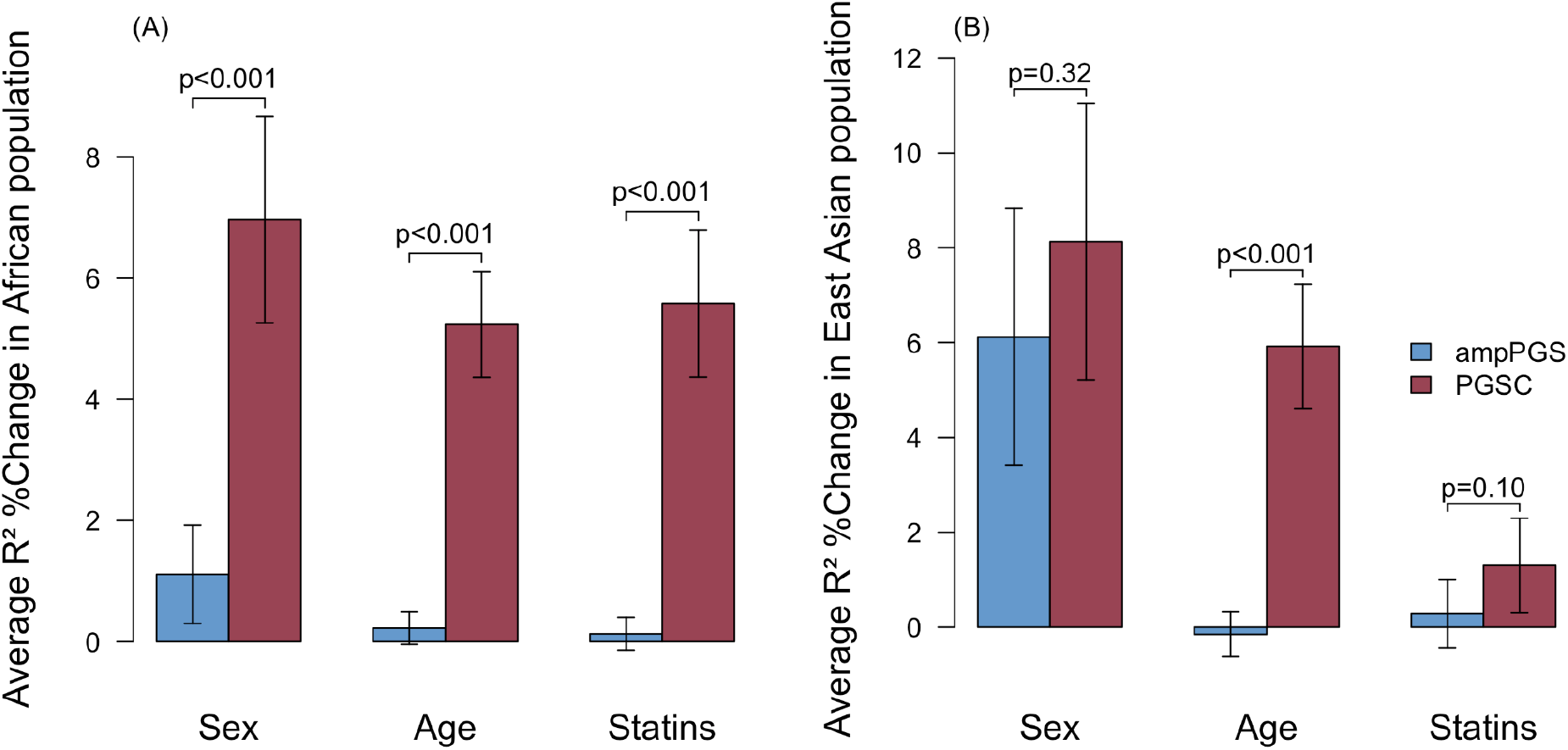
PGSC improves cross-population prediction accuracy in UKB. Average R^2^ %Change for phenotype prediction achieved by PGSC (red) and ampPGS (blue) relative to PGS evaluated in (A) Afr and (B) Asn ancestry individuals from UKB. Phenotypes were included if the PGS R^2^ exceeded 0.01 in both the Euro and the target population (21 phenotypes for Afr, 39 for Asn). Error bars represent 95% bootstrap confidence intervals.

We next evaluated prediction portability, defined as the relative incremental R^2^ in the target population compared to that of the Euro population (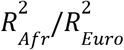 or 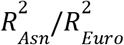). PGSC significantly improved portability for each context and ancestry on average across traits (all 6 p < 0.05, Fig. S6), whereas ampPGS only improved portability using GxSex in the Asn population. Nonetheless, PGSC portability gains were modest (0.26–2.52%) and noisy. Conservatively, we conclude that PGSC improves prediction accuracy across populations but that it does not substantially improve portability *per se*.

### PGSC replication in MSM

GxC signals are notoriously prone to statistical artifacts and often fail to replicate across datasets^19^. To ensure PGSC is learning generalizable signals rather than UKB-specific artifacts, we tested performance in the Mount Sinai Million Health Discovery Program (MSM), a US-based EHR Biobank. We tested 16 phenotypes in 18,493 unrelated individuals genetically similar to European ancestry reference groups (as defined in Belbin et al. 2021; Table S15)^21^, using sex as a context (Methods). PGSC outperformed PGS for 5/16 phenotypes (p < 0.05), significantly improving R^2^ %Change by 7.7% on average (p = 1.4×10^−4^, Fig. 5, Fig. S7). As in UKB, the greatest gain by far was for testosterone (73.6%), with substantial gains for biomarkers such as hemoglobin A1c (13.6%) and triglycerides (9.9%). ampPGS significantly outperformed PGS in 3/16 phenotypes, improving R^2^ %Change by 2.1% on average (p = 1.6×10^−3^). We conclude that performance gains from PGSC replicate across cohorts.

**Fig. 5.**
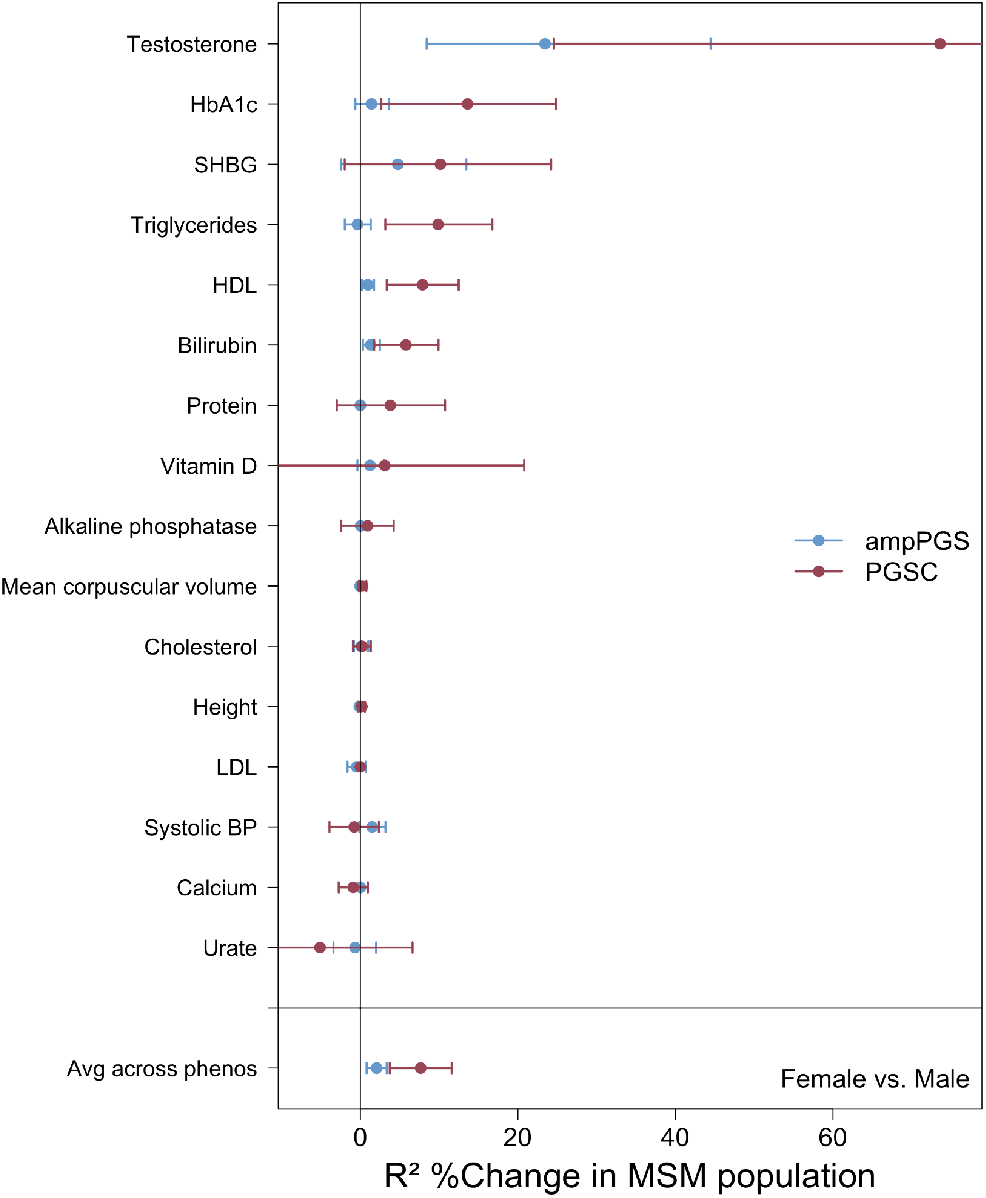
PGSC prediction improvements replicate in MSM. R^2^ %Change for phenotype prediction achieved by PGSC (red) and ampPGS (blue) relative to PGS using sex as a context across 16 traits matched in the MSM Biobank. The x-axis shows the relative change in incremental R^2^; the y-axis lists individual phenotypes, with the average across traits shown at the bottom. Error bars represent 95% confidence intervals from 10,000 bootstrap samples. PGSC, ampPGS, and PGS were trained in unrelated WB individuals from UKB and evaluated in European and Ashkenazi Jewish ancestry individuals in MSM.

### PGSC is partly robust to phenotype scale

Statistically significant GxC signals can appear or disappear depending on how the phenotype is scaled^34,35^. We performed simulations to assess PGSC and ampPGS performance on non-additive phenotype scales. We found that ampPGS and, to a lesser extent, PGSC capture scale-dependent signals that vanish after suitable transformation (Ext Data Fig. 3). This is consistent with PGSC capturing both locus-specific GxC and amplification, with the latter being scale-dependent^17^.

Thus, we asked how our primary analyses in Fig. 3 depend on phenotype scale by re-analyzing 11 phenotypes on the log scale, a simple scale transformation that eliminates many amplification signals in UKB^17^. We found that all significant ampPGS gains with GxSex vanished on the log scale (average R^2^ %Change reduced from 11.6% to 0.2%, Fig. 6). We found that PGSC with GxSex also lost power on the log scale, yet still substantially outperformed PGS (average R^2^ %Change reduced from 13.6% to 7.7%), consistent with simulation results.

**Fig. 6.**
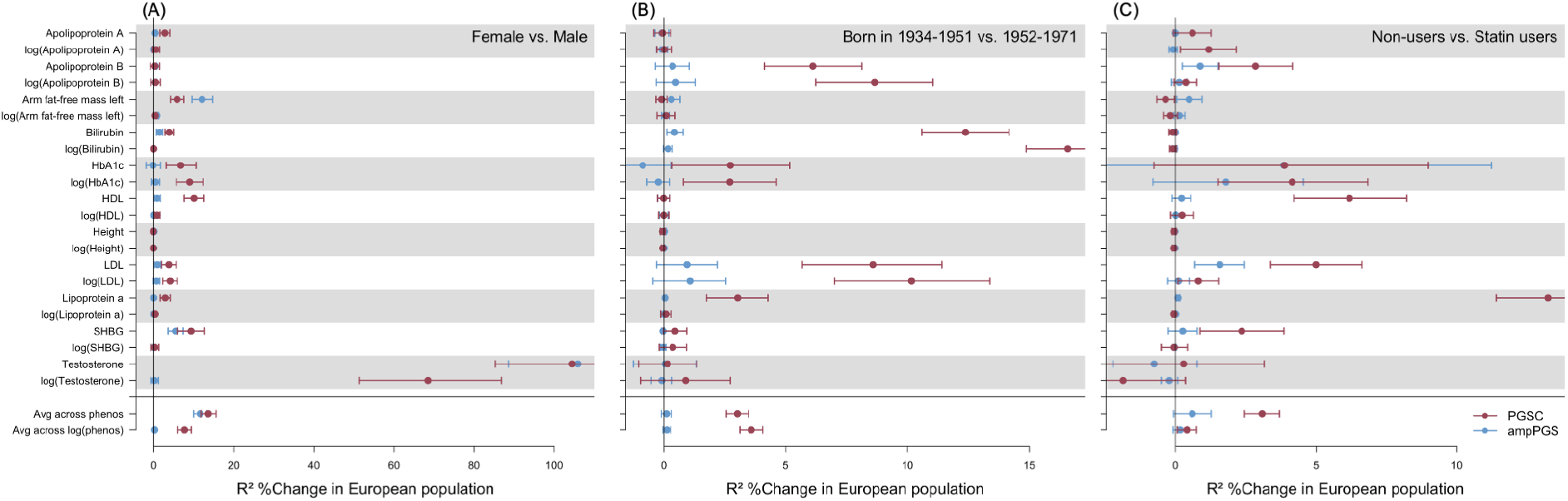
Impact of phenotype measurement scale on PGSC and ampPGS performance. R^2^ %Change for phenotype prediction achieved by PGSC (red) and ampPGS (blue) relative to PGS in the Euro population, shown for both the original and log-transformed phenotype scales. The x-axis indicates the percent improvement in R^2^, and the y-axis lists individual phenotypes with their log-transformed equivalents, as well as the average improvement across phenotypes and log-transformed phenotypes.

Testosterone provides a clear illustration: on the default scale (nmol/L), R^2^ %Change is approximately 100% for both ampPGS and PGSC; on the log scale, it falls to 68.5% for PGSC and 0.2% for ampPGS.

The PGSC GxAge signals, however, were generally preserved or even strengthened by log transformation (R^2^ %Change increased from 3.0% to 3.6%). In particular, this suggests that the default-scale heteroskedasticity signals captured by PGSC (Ext Data Fig. 1) likely reflect a genuine increase in nongenetic variation with age. Lp(a) is a clear exception, where PGSC R^2^ %Change drops from 3.0% to 0.1%, highlighting how scale-dependent GxC varies across traits and contexts.

For GxStatins, PGSC gains for Lp(a), ApoB, LDL, and HDL vanished under log transformation. By contrast, the signals for ApoA and HbA1c increased (R^2^ %Change from 0.6 to 1.2% and 3.9 to 4.1%, respectively). The strengthened HbA1c result is consistent with our prior results indicating that distinct genetic effects act on HbA1c in statin users and non-users. In contrast, genetic effects on cholesterol are more consistent with genome-wide buffering by statins^4^.

Together, these results show how GxC results from a mix of log-additive amplification effects and scale-independent locus-specific effects, where ampPGS captures the former, and PGSC captures both.

## Discussion

We introduced PGSC, a framework to incorporate locus-specific GxC effects into PGS. PGSC can provide substantial improvements over standard PGS when GxC heritability is high, such as GxSex effects on testosterone (104.5%) or GxAge effects on cholesterol (7.6–8.6%). PGSC is robust because it reliably simplifies to standard PGS when GxC heritability is low. In contrast to most studies of GxC and PGS, we demonstrated that PGSC predictions replicated out-of-sample, across ancestries, in an independent cohort, and for many phenotypes across three contexts. The genome-wide GxC model, ampPGS, outperformed PGS for a smaller set of phenotypes, primarily for GxSex. Together, our results directly demonstrate that locus-specific GxC effects can robustly improve polygenic prediction.

Our evaluation of log-scale phenotypes revealed an underappreciated issue in testing non-additive PGS: large gains may vanish entirely on another scale. We found that log transformation eliminated every case where ampPGS outperformed PGS, as expected if genetic effects and context are primarily log-additive^17^. Conversely, log transformation only partly reduced PGSC gains, because PGSC captures both scale-dependent amplification and scale-independent locus-specific GxC.

Our results deliver on several lines of evidence suggesting that GxC should improve PGS prediction. First, studies have shown PGS do not transfer well across ancestries^13,14^, age^12^, or treatment status^3,4,25^. Theoretically, these transferability gaps could be explained by unmodeled GxC, as PGS are implicitly optimized for the contexts in the training data rather than the target individuals. Second, variance component models have demonstrated that GxC effects can explain substantial heritability, which standard PGS cannot capture^5–10^. Similar arguments have been made on a per-variant basis^23^ or by testing in-sample interaction between a context and an additive PGS^26^ or a GxC-based PGS^36,37^. Third, a Bayesian GxSex PGS model was found to improve cross-validated prediction accuracy^10^, but this is confounded by the benefits of Bayesian shrinkage of additive effects^38^ (Table S1).

PGSC has potential utility for precision medicine by estimating an individual’s treatment response as PGSC(treated) - PGSC(untreated). We recently identified GxStatins effects on LDL—the primary target of statins—and HbA1c—a potential side effect of statins^39–41^—at the level of individual genes, cell types, PGS, and genome-wide heritability^4,8,42^. Here, we show that incorporating GxStatins can improve their cross-sectional polygenic prediction. The HbA1c result persisted on the log scale, while cholesterol-related results did not, suggesting the former interactions may reflect heterogeneous biological responses while the latter more likely reflect genome-wide buffering. In the future, it will be essential to validate these predictions prospectively, given the endogeneity inherent in observational cohorts^3,4^. Nonetheless, cross-sectional approaches are needed to predict polygenic treatment response, as GWAS-scale randomized controlled trials are infeasible.

PGSC significantly improved prediction in Afr and Asn UKB populations (5.40% on average), but only modestly improved portability (0.26–2.52%), i.e., the prediction R^2^ in each target population relative to the Euro UKB population. This suggests that GxC is not a major limitation to cross-population portability, consistent with recent studies that primarily implicate MAF and LD differences instead^43–48^. In turn, this suggests that GxC models cannot close the portability gap, and instead that more diverse data collection is needed. Nonetheless, we only studied cross-population portability within UKB, so GxC could play a larger role when porting across biobanks^49^, countries, or healthcare systems. Relatedly, PGSC is likely to be useful for recently admixed populations by modeling each individual’s admixture proportions as a context rather than the standard approach that bins them all together, e.g., “Hispanic or Latino^50^.”

Our approach has limitations. First, PGSC uses binarized context measures for simplicity and interpretability. Extending to continuous contexts is theoretically straightforward but requires further consideration for robust extrapolation, such as appropriate context scaling; related concerns apply to extending PGSC to binary disease traits^51^. Second, PGSC uses univariate context measures, prohibiting inclusion of multi-level contexts (e.g., never-, current-, or previous-smoker) or disentangling correlated contexts (e.g., GxStatins and GxAge). Nonetheless, PGSC results could be combined across contexts *post hoc* to further improve prediction. Finally, we used standard clumping and thresholding (C+T) as the underlying tuning method in PGS, PGSC, and ampPGS, which provides a fair like-with-like evaluation, but more sophisticated models^10,52–54^ should further improve accuracy for all approaches.

Overall, PGSC demonstrates that GxC interactions can rigorously improve polygenic prediction accuracy, a key step toward context-aware clinical prediction of disease risk and treatment response.

## Methods

### PGSC model

PGSC is built in two stages (Note S3). First, GWAS and GxC GWAS are performed to obtain SNP additive effect sizes (β) and GxC effect sizes (λ). These GWAS use covariates *X* (e.g., genetic PCs), a binarized context vector *C*, and a per-SNP genotype vector *G*, regressed on a phenotype *y*:

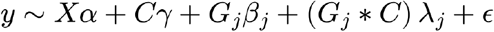

The context *C* is defined as:

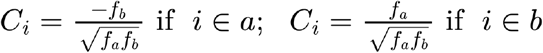

where *f*_*a*_ and *f*_*b*_ are the fraction of individuals in context groups *a* and *b*, respectively, so that *C* has mean zero and variance one. The GxC term (*G*_*j*_ * *C*) represents element-wise multiplication of genotype *G*_*j*_ and context *C*, which is orthogonal to *G*_*j*_—assuming *G*_*j*_ has equal variance in both contexts. Residuals (ϵ) are assumed i.i.d. Gaussian.

Second, PGSC combines the additive and GxC effect estimates into a polygenic score. The additive PGS is constructed using standard clumping and thresholding (C+T), where *G* is the genotype matrix, 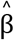 is the vector of additive GWAS effect estimates, and *τ* is the p-value inclusion threshold:

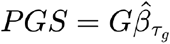

The GxC term 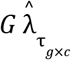 adds a second component built from GxC GWAS effect estimates 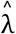, parameterized by its own p-value threshold *τ*, and context-specific tuning parameters ρ_*a*_ and ρ_*b*_ :

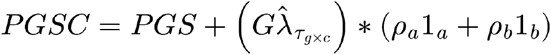

where 1_*a*_ and 1_*b*_ are indicator vectors for contexts *a* and *b*.

Both the additive and GxC polygenic terms are constructed using C+T with a fixed clumping threshold of *R*^2^ = 0. 1 and a 250kb window. All tuning parameters (*τ*_*g*_, *τ*_*g*×*c*_, ρ_*a*_, ρ_*b*_) are optimized to maximize prediction accuracy in a tuning cohort. *τ*_*g*_ is chosen first to define the PGS; *τ*_*g*×*c*_, ρ_*a*_ and ρ_*b*_ are then jointly optimized given the fixed PGS. *τ*_*g*_ and *τ*_*g*×*c*_ are each selected from a fixed grid of p-value thresholds (10^−10^, 10^−8^, 10^−6^, 10^−4^, 0. 001, 0. 005, 0. 01, 0. 05, 0. 1, 0. 5); for each value of *τ*_*g*×*c*_, the optimal ρ_*a*_ and ρ_*b*_ are obtained by least squares.

Under the homoskedastic additive model, PGSC approximately reduces to the standard PGS by learning ρ_*a*_ ≈ ρ_*b*_ ≈ 0, giving the GxC term nearly zero weight.

### ampPGS model

We compared PGSC to a simpler GxC model, ampPGS, defined as:

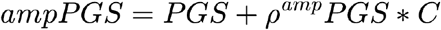

where ρ^*amp*^ is a tuning parameter optimized in the tuning cohort. When ρ^*amp*^ > 0, the PGS effect is amplified for individuals in the positive-encoded context; when ρ^*amp*^ = 0, ampPGS reduces to the PGS.

ampPGS assumes that context uniformly scales all SNP effects genome-wide, implying that GxC effects are proportional to additive effects (λ = *t*β for scalar *t*). PGSC is strictly more general: it reduces to ampPGS when this proportionality holds exactly, and to standard PGS when *t* = 0. Many prior studies have used this model to detect polygenic GxC by testing if ρ^*amp*^ = 0^11^; here, we use this model to perform out-of-sample prediction.

### GWAS and GxC GWAS

GWAS and GxC GWAS were performed in the UKB White British (WB) training population using PLINK 2.0^22^ (v2.00a6LM AVX2 Intel, 4 Jul 2024) via the --glm flag, with the interaction modifier added to include GxC interaction terms in the GxC GWAS. Each binarized context was included as a covariate in both the GWAS and GxC GWAS.

### Score construction and tuning using PLINK 2.0

Clumping was performed in the WB training population using PLINK 2.0^22^ with a 250kb window and *R*^2^ = 0. 1, including all SNPs at *p* ≤ 1 in the clumping procedure. We then computed additive PGS and GxC PGS profiles across all p-value threshold combinations in the tuning cohort using --score, applying the clumped SNP list with additive 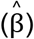 and 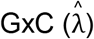 effect estimates, respectively. We selected optimal thresholds (*τ*_*g*_, *τ*_*g*×*c*_) and tuning parameters (ρ_*a*_, ρ_*b*_) by maximizing incremental *R*^2^ over standard PGS in the tuning cohort. We generated final PGSC predictionsin the held-out validation cohort using --score, leveraging the optimized parameters.

### Measuring and comparing prediction accuracy

We evaluated prediction accuracy using incremental Pearson *R*^2^ conditional on the variance explained by covariates. To compare prediction accuracy across models, we computed the absolute change in incremental *R*^2^:

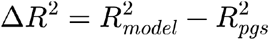

and plotted the Δ*R*^2^%, defined as 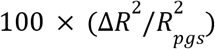, where 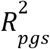 is the incremental *R*^2^ of the baseline model, typically the standard additive PGS, and 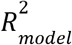 is the incremental *R*^2^ of the model under evaluation (e.g., PGSC or ampPGS). We assessed statistical significance of Δ*R*^2^ using a paired bootstrap test with 10,000 bootstrap (*b*) samples. Individuals were resampled with replacement, and Δ*R*^2^ was recomputed:

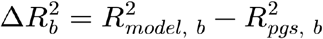

This paired resampling approach accounts for the correlation between scores evaluated in the same individuals, substantially improving power relative to unpaired tests assuming independence.

We summarized average performance across traits as the unweighted mean of Δ*R*^2^ % across included phenotypes. We computed the standard error of the mean assuming phenotypes were independent.

### Simulations

We constructed a standard quantitative genetics simulation with additive effects, locus-specific GxC, genome-wide amplification GxC, and heteroskedasticity (Note S2). In brief, we simulated *S* = 10, 000 independent binomial SNPs with minor allele frequencies drawn uniformly from (0.05, 0.5), and randomly selected *S*_*caus*_ causal SNPs. At each causal locus, we drew additive and locus-specific GxC effects independently from Gaussian distributions, scaled so that each explained a prescribed fraction of phenotypic variance. Context *C* represented a binary variable, and the phenotype equaled the sum of additive, GxC, context-main, and residual effects.

Amplification GxC denoted interaction effects proportional to the additive effects; locus-specific GxC denoted interaction effects drawn independently of additive effects. Residual noise could be made heteroskedastic across context groups, with the residual variance in one context group set to η times that of the other. This framework is general and allowed us to vary the number of individuals, causal SNPs, and the phenotypic variance explained by additive effects, locus-specific GxC, amplification GxC, and context main effects, as well as the context imbalance and heteroskedasticity.

Each simulation used three independent, equally sized populations. Additive (GWAS) and interaction (GxC GWAS) effects were estimated in the training population; the additive p-value threshold (*τ*_*g*_), the GxC p-value threshold (*τ*_*g*×*c*_), and the tuning parameters (ρ) were learned in the testing population by maximizing prediction R^2^; out-of-sample R^2^ was reported using the validation population. We selected both *τ*_*g*_ and *τ*_*g*×*c*_ over a fixed grid of p-value thresholds (10^−10^, 10^−9^, 10^−8^, 10^−7^, 10^−6^, 10^−5^, 10^−4^, 0. 001, 0. 01, 0. 1, 1

Finally, we simulated phenotype scale transformations by simulating phenotypes as above and then applying a Box-Cox power transformation with coefficient λ_*BoxCox*_ (Note S2.2), swept over [−1, 2]. As in Costantino et al. 2026^17^, we set *y* to *y* − *min*(*y*) + 1 to ensure it was positive before applying the transformation, and then re-standardized to mean zero and variance one.

### UKB data

The UKB is a large dataset of genotypic and phenotypic data collected from approximately 500,000 United Kingdom residents recruited between 2006 and 2010 at ages 40–69^20^.

In this study, we included 370,357 individuals who passed UKB quality control measures. We excluded individuals related at the third degree or closer, as defined by UKB data field 22020, and all individuals who had withdrawn from UKB before analysis. Remaining individuals were assigned to one of four ancestry groups — White British (WB), European (Euro), African (Afr), and East Asian (Asn) — and individuals not assignable to these groups were excluded. UKB defines WB individuals as those who self-identify as White British and cluster tightly in genetic principal component (PC) space^20^. The Euro group comprises individuals self-reporting as “All other White” or “Irish” based on UKB self-report data. We defined Afr and Asn ancestry groups following Dahl et al. 2023^28^.

Genotype QC was applied per ancestry group according to its intended use in the analysis. For the WB training population, SNPs were filtered for missing call rate > 0. 01 (PLINK 2.0^22^; --geno), minor allele frequency (MAF) *<* 0. 01 (--maf), and Hardy-Weinberg equilibrium (HWE) exact test *p <* 10^−6^ (--hwe), yielding a final set of 6,529,173 SNPs. We filtered individuals across all four ancestry groups for missing call rate > 0. 01 (--mind). Final sample sizes after QC are provided in Table S3.

#### Covariates

We performed all GWAS using PLINK 2.0^22^, adjusting for age (derived from year of birth), sex, and the first 10 genetic PCs as provided by UKB (computed across all participants). We also included each binarized context (sex, age, and statin usage) as a covariate. In analyses where sex or age were used as contexts, we replaced the continuous UKB-provided values with their binarized equivalents to avoid collinearity.

#### Contexts

We included 3 contexts in this study: sex, age, and statin usage. We coded sex as female or male based on UKB sex (field 31). We binarized age at the median year of birth (field 34) in the dataset, coding individuals born before the median as older and those born at or after the median as younger; the median birth year was 1952. We defined statin usage following Sadowski et al. 2024^4^ and coded individuals as statin users or non-users based on self-reported medication data.

#### Phenotypes

We analyzed 53 quantitative phenotypes in total: 52 were obtained directly from UKB using the most recent available instance measurement, and we constructed waist-to-hip ratio adjusted for BMI as defined in Zhu et al. 2023^10^. UKB field codes for all phenotypes are provided in Table S2. Traits were included in each evaluation for a given population if the baseline PGS *R*^2^ exceeded 0. 01 in both that target population as well as the Euro population, applied separately for each ancestry group (Fig. S8).

### MSM replication cohort

The MSM Biobank consists of electronic health records and genetic data from approximately 60,000 participants in the Mount Sinai Health System in New York. Participant recruitment occurred between 2007 and 2023. This study was approved by the Icahn School of Medicine at Mount Sinai’s Institutional Review Board (IRB 07–0529). All study participants provided written informed consent.

MSM participants were genotyped using the Illumina Infinium Global Diversity Array (GDA; number of participants, N=23,430; number of variants, n=1,833,111) or Infinium Global Screening Array (GSA; N=32,595; n=635,623). Quality control consisted of removing participants with a call rate < 95%, a mismatch between self-reported and genetic sex, and/or high heterozygosity. Duplicated sites and sites with a genotyping rate of < 95% were removed. QC was performed using PLINK 2.0^22^. We imputed data with the TOPMed imputation server^55^ using the hg38 genome build. We subsetted imputed sites to the positions included in the UKB, for a total of 6,196,203 overlapping SNPs. We calculated 10 PCs across participants using PLINK 2.0 for the genotyping sites after LD pruning.

We used identity-by-descent clustering, a validated approach for assigning fine-scale ancestry^21^, to assign MSM participant ancestry. Briefly, we called pairwise identity-by-descent using iLASH^56^, and performed unsupervised Louvain clustering^57,58^ implemented in the Python package NetworkX^59^. Clusters represented groups of people who shared more IBD with participants within the cluster than with those outside it. We assigned cluster labels using reference data from the 1000 Genomes Project, HGDP, and SGDP, along with self-reported questionnaire data. In this study, we considered one cluster with predominant European ancestry (n=18,493), representing patients with Ashkenazi Jewish, Italian, Irish, and North European genetic ancestries.

We processed MSM laboratory data according to the QualityLab pipeline^60^. Briefly, we cleaned quantitative lab values obtained in different clinical contexts (ambulatory, emergency, inpatient, and urgent care). We considered only labs with at least 100 patients and at least 1000 numeric observations. Only adult (> 18 years) lab values were considered. For each lab, we removed outliers, defined as values greater than or less than 4 standard deviations from the lab mean. Non-numeric labs were removed. Lab tests measuring the same analyte were combined when > 70% of the reporting units matched. After cleaning, we calculated a median value and median age per person for each lab test in each clinical context and across all contexts. We manually matched laboratory phenotypes to UKB phenotypes using available metadata. Sex was self-reported sex indicated by the participant in questionnaire data. We calculated height as the median value across visits recorded in the electronic health record. The full set of individual counts per phenotype is in Table S15. We initially applied standard PGS, ampPGS, and PGSC across 36 phenotypes, retaining 16 after further QC based on thresholding for PGS *R*^2^ > 0. 01 in MSM (Fig. S9) and sample size > 500 individuals per trait (Table S15).

## Extended data figures

**Extended Data Fig. 1.**
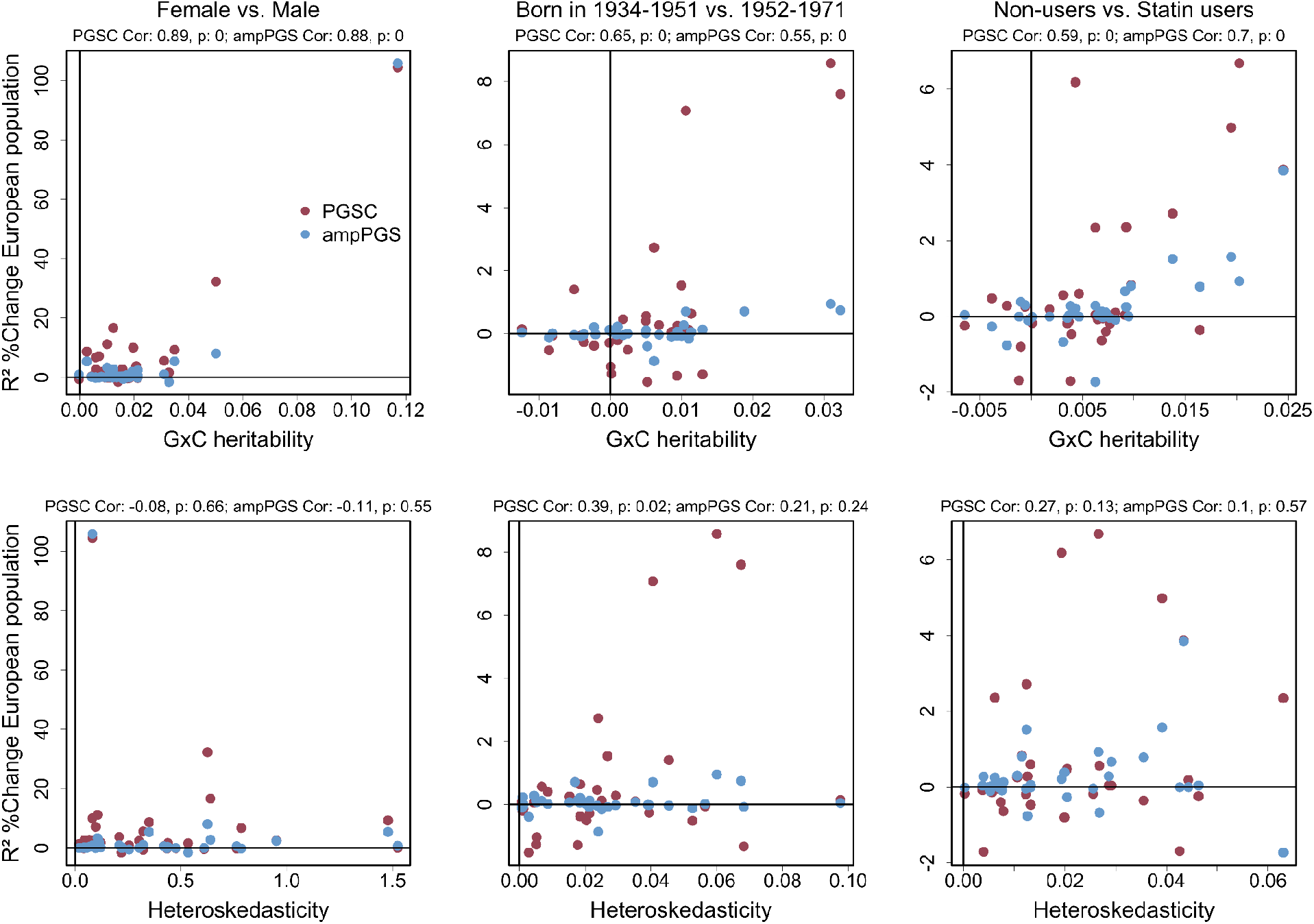
Genetic architecture underlying PGSC prediction improvements in UKB. Each point is one phenotype. Top row: GxC heritability estimated by GENIE versus R^2^ %Change for PGSC (red) and ampPGS (blue) relative to standard PGS, for contexts sex, age, and statins. Bottom row: Context-dependent heteroskedasticity estimated by GENIE plotted against R^2^ %Change for PGSC and ampPGS per context. Pearson correlations and associated p-values for each method are shown above each panel. Analysis is restricted to the 33 UKB phenotypes present in both the PGSC and GENIE analyses.

**Extended Data Fig. 2.**
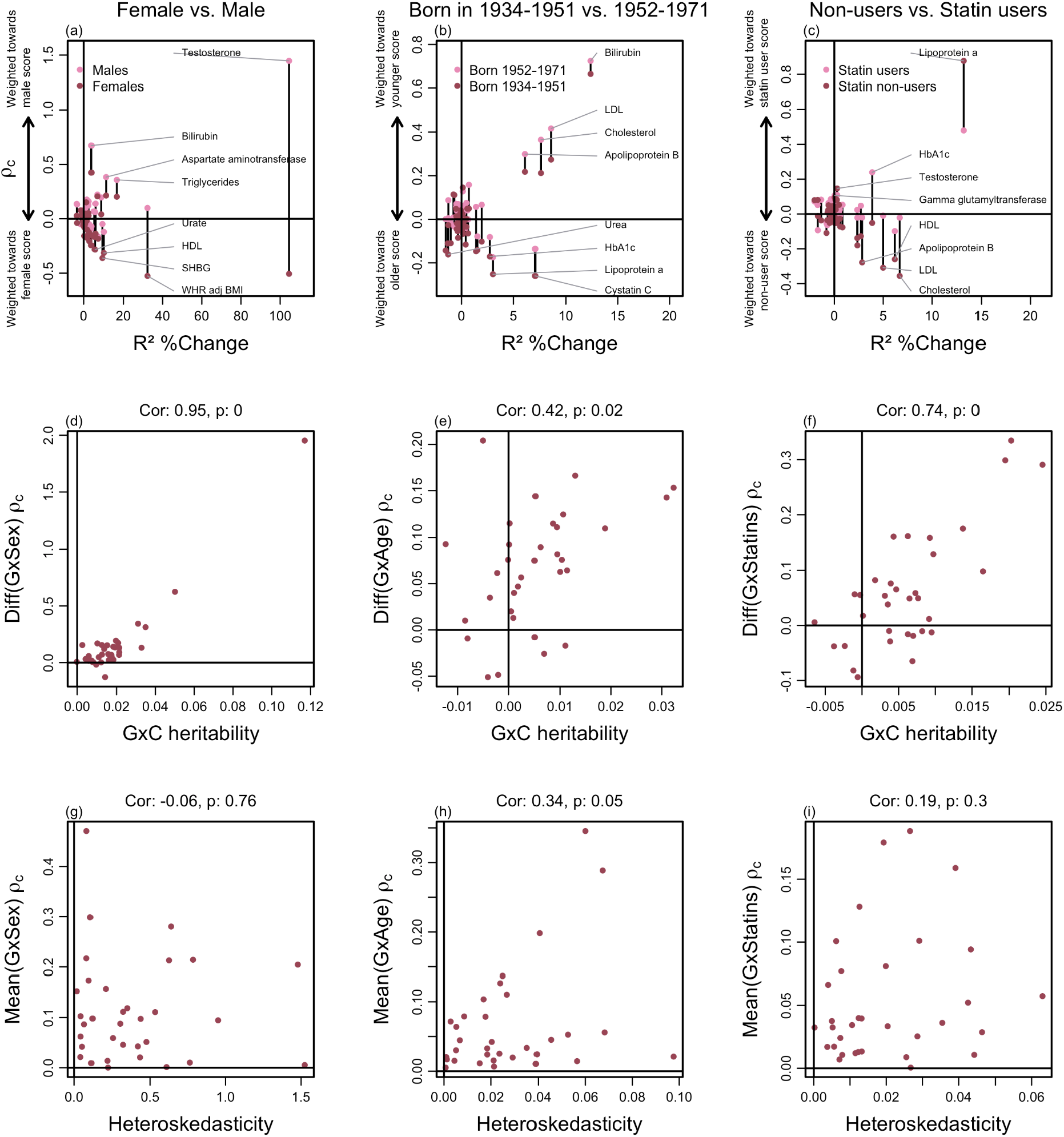
Tuning parameters underlying PGSC prediction improvements in UKB. (a-c) PGSC tuning parameters (ρ) plotted against R^2^ %Change for phenotype prediction relative to PGS, for contexts of sex, age, and statin treatment status. Each point represents a single phenotype evaluated in the Euro UKB population. For each context, the two ρ_*c*_ are shown connected by a black line: ρ_*male*_, ρ_*younger*_, and ρ_*statins*−*users*_ (pink), and ρ_*female*_, ρ_*older*_, and ρ (red). Larger absolute ρ_*non*−*users*_ values indicate greater weighting of the GxC PGS term for the corresponding context group, as shown by the arrows to the left of each panel. Larger distance between ρ_*c*_ values indicates increasing 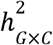, and larger distance between the mean ρ_*c*_ value per phenotype and zero indicates increasing heteroskedasticity (η). The four phenotypes with the largest and smallest ρ_*c*_ values are labeled in each panel. (d-f) Difference in PGSC tuning parameters between the two context groups plotted against genome-wide GxC heritability estimated independently with GENIE. Pearson correlation coefficient and p-value are reported in each panel. (g-i) Absolute value of the mean PGSC tuning parameter across the two context groups plotted against absolute genome-wide heteroskedasticity estimated with GENIE. Pearson correlation coefficient and p-value are reported in each panel. The 33 UKB phenotypes overlapping those studied in Pazokitoroudi et al. 2024^8^ are shown in d-i.

**Extended Data Fig. 3.**
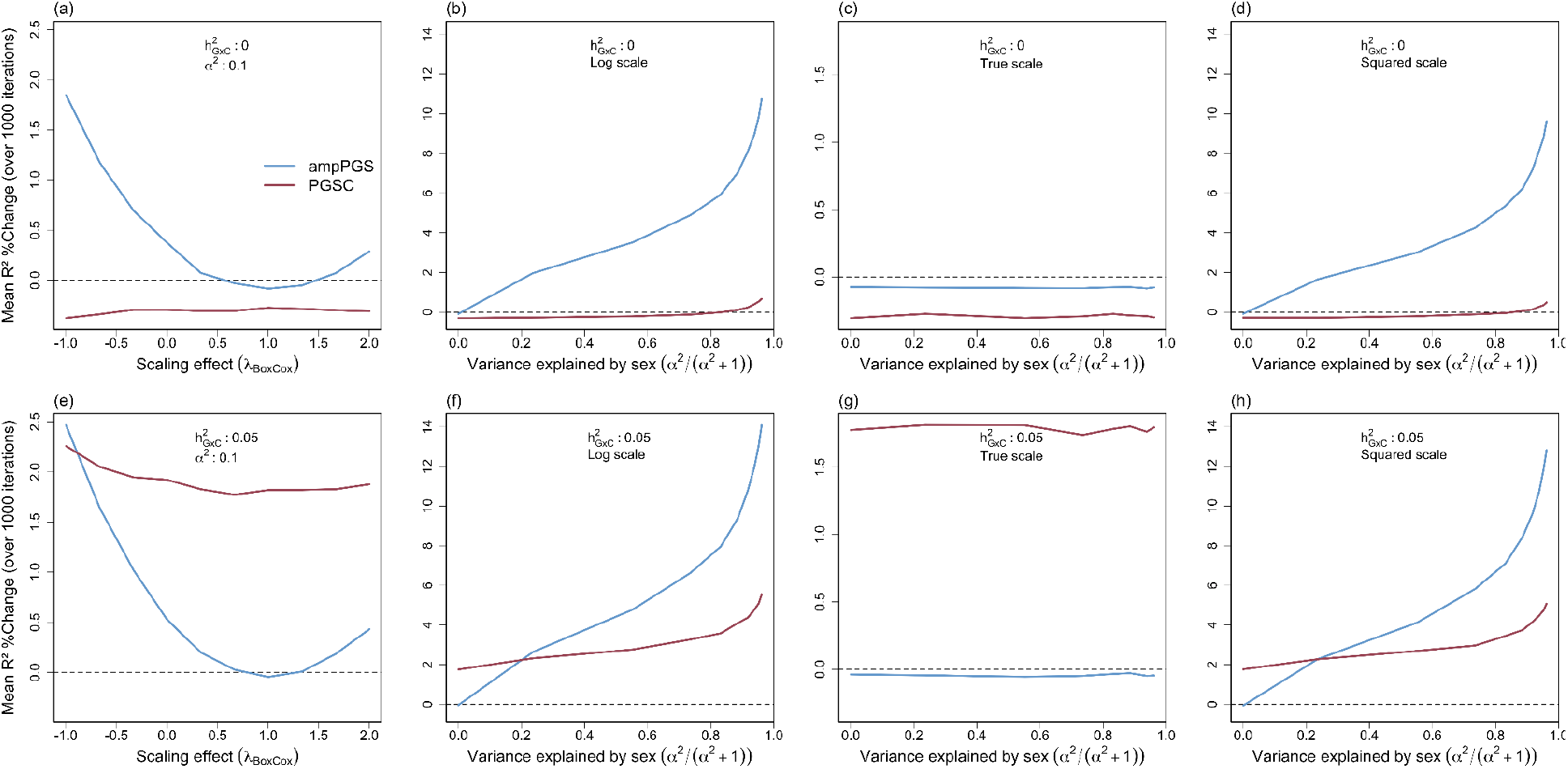
Effect of phenotype scaling and variance explained by context across scales on PGSC and ampPGS. R^2^ %Change for phenotype prediction achieved by PGSC (red) and ampPGS (blue) relative to PGS across 1,000 simulation replicates with locus-specific GxC. Simulation parameters are: 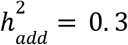; *S* = 10, 000; *S*_*caus*_ = 1,000; *N* = 10, 000; *C*_*prob*_ = 0. 5; 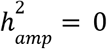; (a-d) use 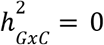, and (e-h) use 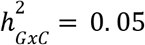. (a,e) sweep the Box-Cox scale from −1 to 2 with the main effect of context fixed at *α*^2^ = 0. 1. (b-d,f-h) sweep along *α*^2^/(*α*^2^ + 1) while λ_*BoxCox*_ is set to either 0 (log scale), 1 (true scale), or 2 (squared scale). We fix the combined variance of genetics and residual noise at 1 and let context add variance *α*^2^ (a-h), so that the fraction of total phenotypic variance explained by context is *α*^2^/(*α*^2^ + 1) on the untransformed scale.

## Author contributions

R.F. and A.D. developed the statistical methodology, conducted the analyses, and wrote the manuscript. C.C. and E.E.K. conducted MSM analyses. M.C. and O.D. contributed to UKB analyses.

## Code availability

Code to implement PGSC is available at https://github.com/reneemf/PGSC. Code to reproduce results is available at https://github.com/reneemf/PGSC_figures.

## Acknowledgments

This research has been conducted using the UK Biobank Resource under Application Number 89052. We are grateful to the UKB and MSM participants, and their families. This work was funded by R35GM150822 (to A.D.). R.F. is funded by the Ford Foundation. M.C. is funded by the Fonds de Recherche du Québec Santé. Compute resources for this study were provided by the Randi HPC Cluster maintained by the Center for Research Informatics (CRI) at the University of Chicago. The Center for Research Informatics is funded by the Biological Sciences Division and the Institute for Translational Medicine/CTSA (NIH UL1TR002389) at the University of Chicago. This work was supported in part through the computational and data resources and staff expertise provided by Scientific Computing and Data at the Icahn School of Medicine at Mount Sinai and supported by the Clinical and Translational Science Awards (CTSA) grant UL1TR004419 from the National Center for Advancing Translational Sciences. Research reported in this publication was also supported by the Office of Research Infrastructure of the National Institutes of Health under award numbers S10OD026880 and S10OD030463. The content is solely the responsibility of the authors and does not necessarily represent the official views of the National Institutes of Health.

## Supplementary figures

**Fig. S1.**
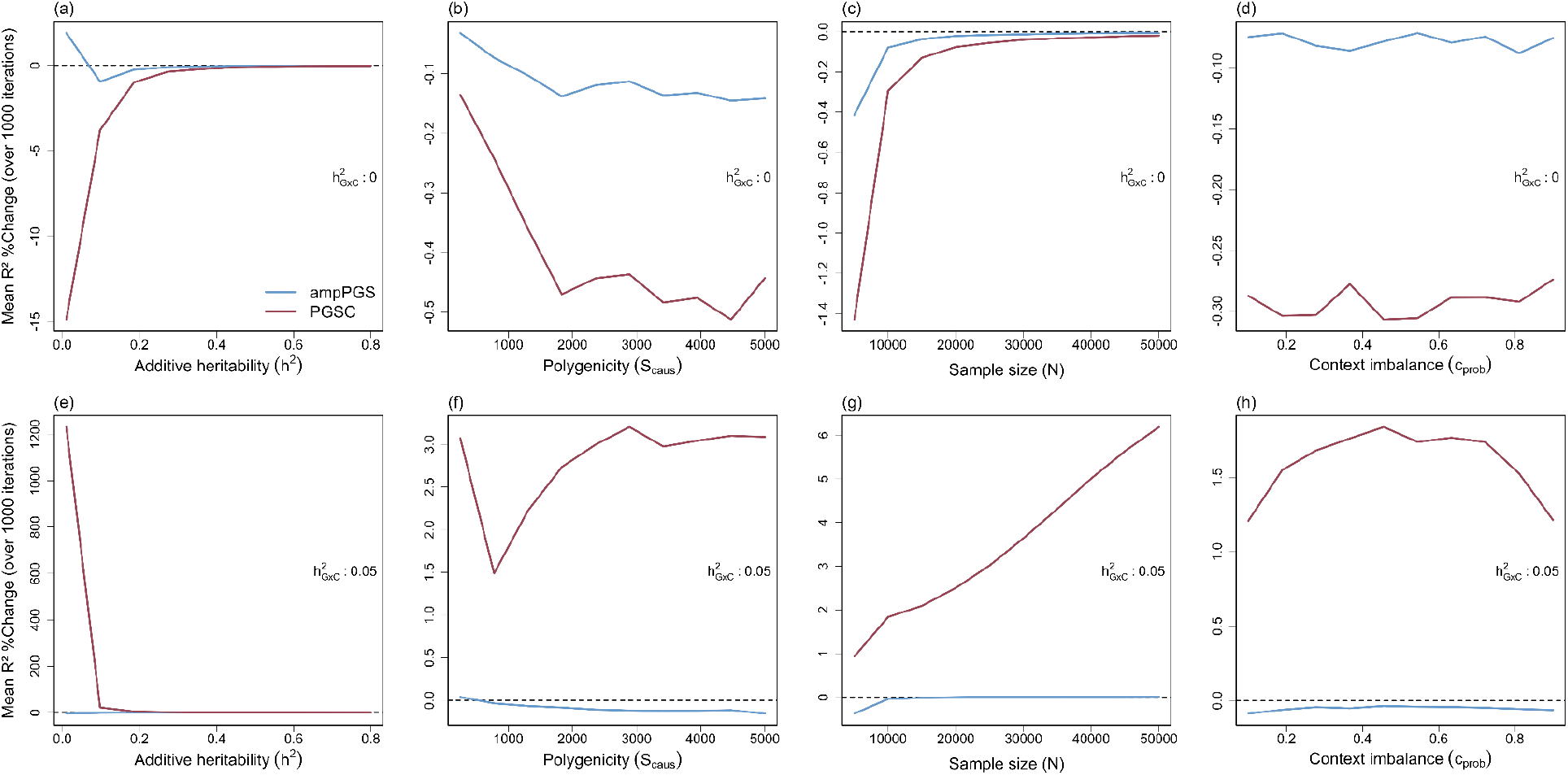
PGSC and ampPGS sensitivity to genetic architecture and study design. R^2^ %Change for phenotype prediction achieved by PGSC (red) and ampPGS (blue) relative to PGS across 1, 000 simulated iterations, given two base locus-specific GxC effects, (a-d) 0 or (e-h) 0. 05. Each panel sweeps one parameter while holding the others at their defaults (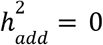. 3; *S* = 10, 000; *S* _*caus*_= 1, 000; *N* = 10, 000; *C*_*prob*_= 0. 5; 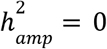).

**Fig. S2.**
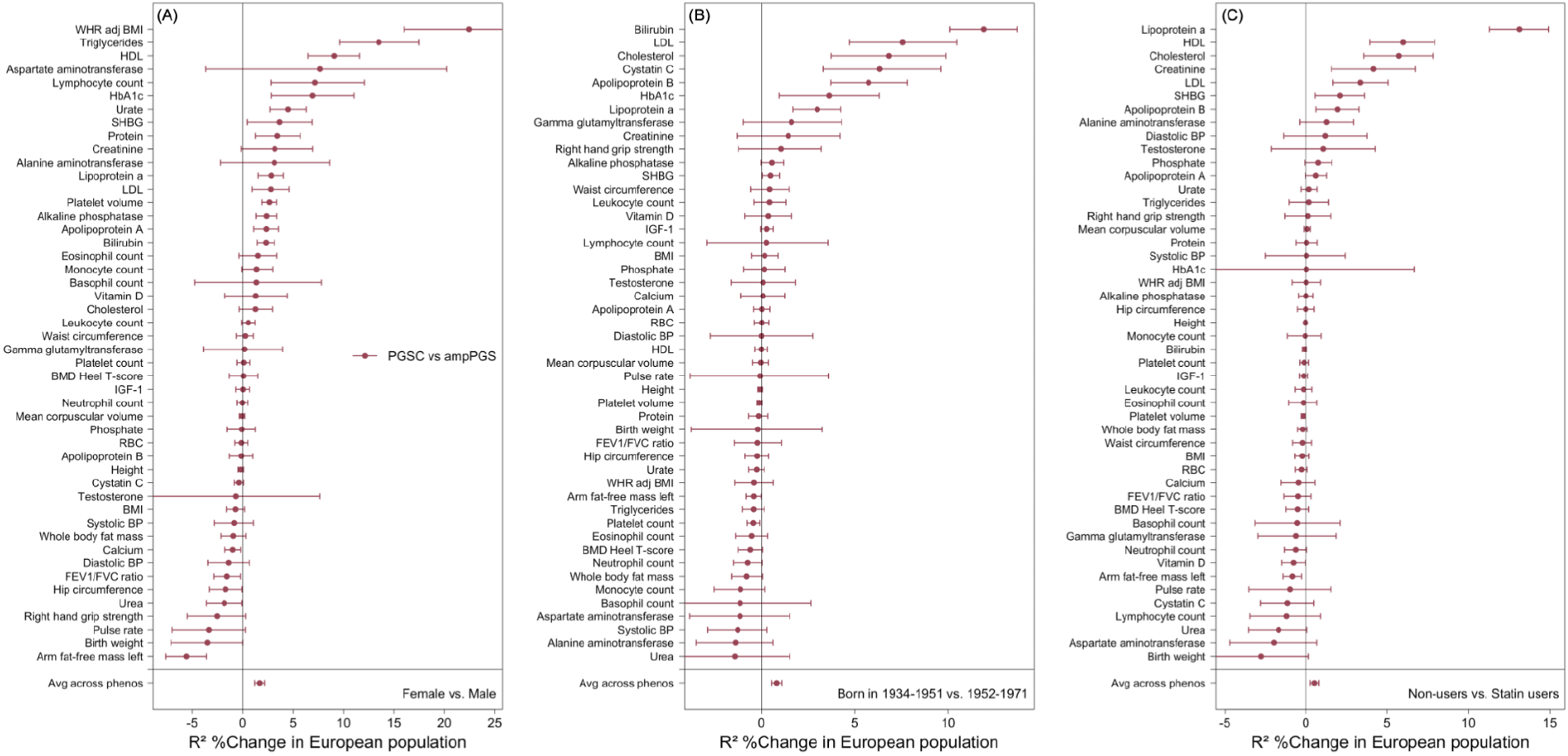
PGSC improves phenotype prediction across contexts compared to ampPGS in the UKB European population. R^2^ %Change for phenotype prediction achieved by PGSC (red) relative to ampPGS across 48 quantitative traits, for contexts (A) sex, (B) age, and (C) statin treatment status. The x-axis shows the relative change in incremental R^2^, while the y-axis lists individual phenotypes, with the average improvement across all traits shown at the bottom. Error bars represent 95% confidence intervals computed from 10,000 bootstrap samples. All scores were trained in a set of unrelated WB individuals and evaluated in an independent set of unrelated European ancestry (Euro) individuals from the UKB.

**Fig. S3.**
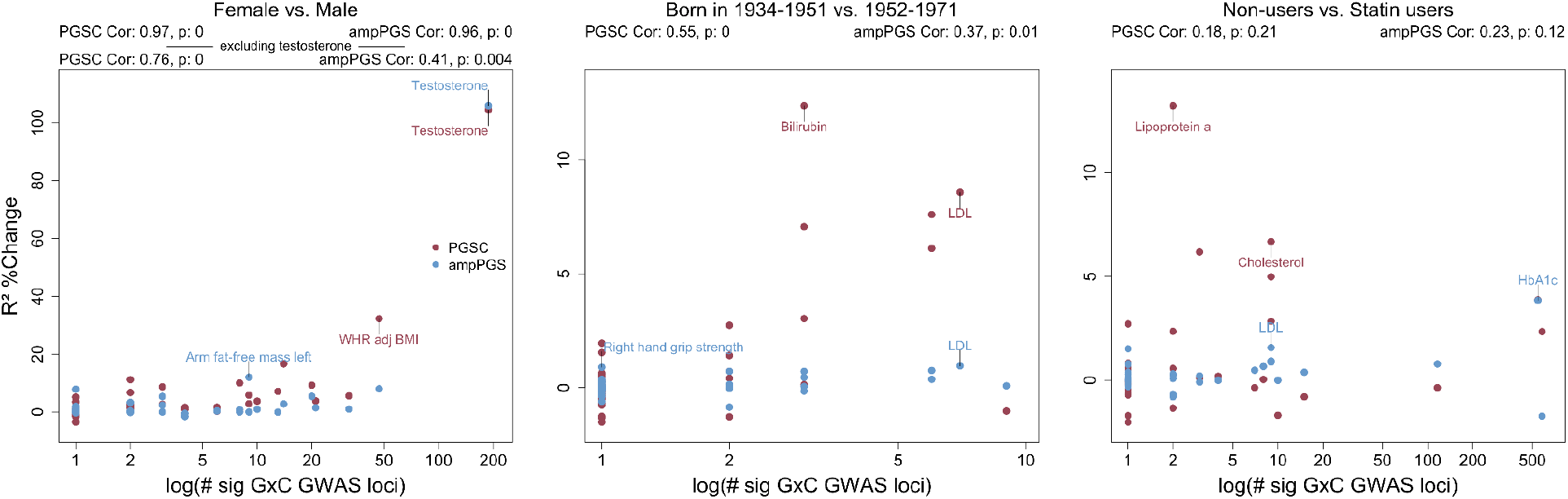
log(#sig GxC GWAS loci) vs R^2^ %Change in UKB Euro population. Each panel shows the R^2^ %Change for the PGSC (red) and ampPGS (blue) in the Euro population relative to the baseline PGS, plotted against the number of genome-wide significant GxC loci per phenotype, across three contexts: sex, age, and statin use. The two phenotypes with the largest R^2^ %Change for each model are labeled. Text above each panel reports the Pearson correlation coefficient and associated two-sided p-value.

**Fig. S4.**
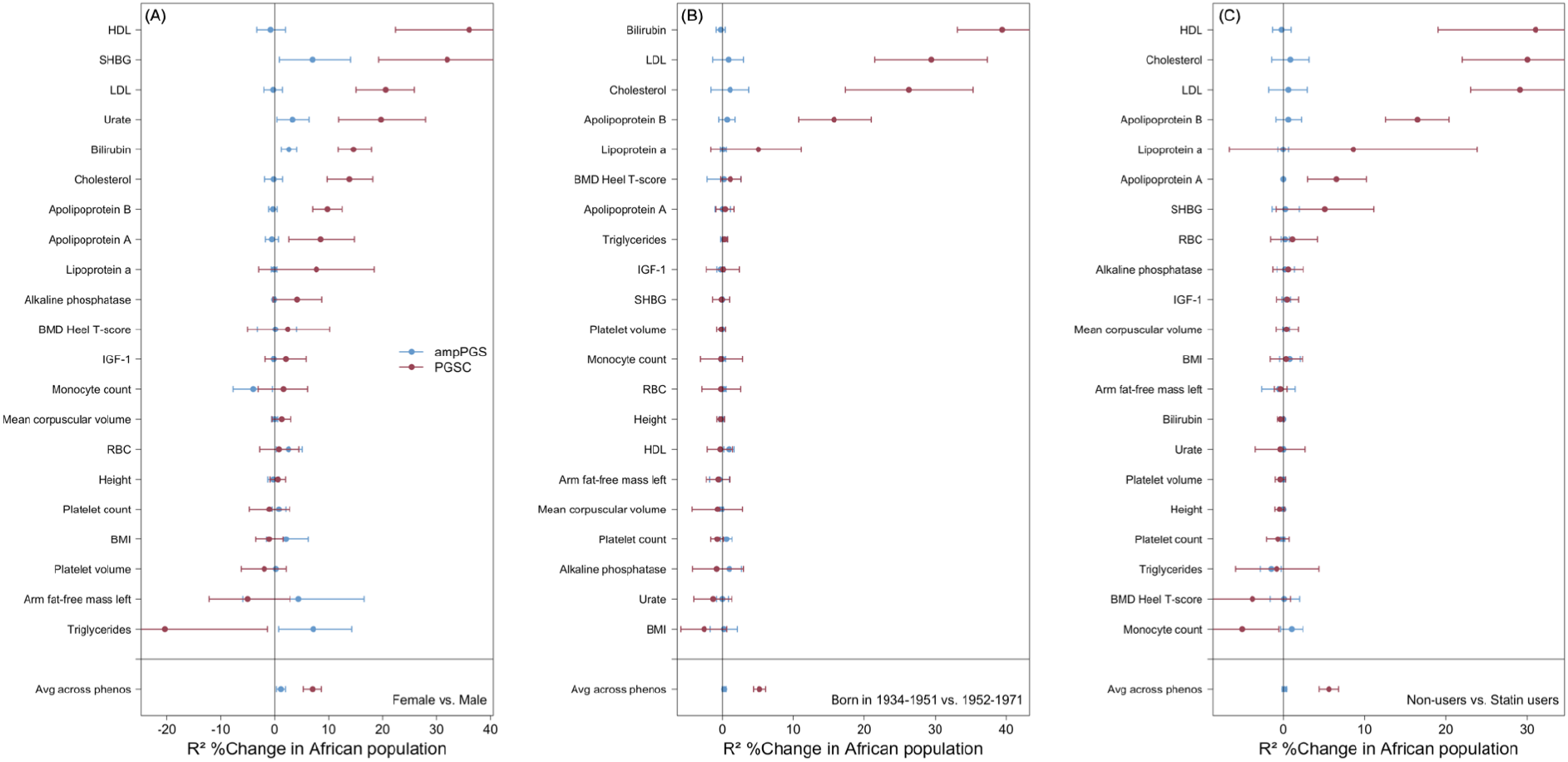
PGSC improves phenotype prediction across contexts in the UKB African population. R^2^ %Change for phenotype prediction achieved by PGSC (red) and ampPGS (blue) relative to PGS across 21 quantitative traits, for contexts (A) sex, (B) age, and (C) statin treatment status. The x-axis shows the relative change in incremental R^2^, while the y-axis lists individual phenotypes, with the average improvement across all traits shown at the bottom. Error bars represent 95% confidence intervals computed from 10,000 bootstrap samples. All scores were trained on a set of unrelated WB individuals and evaluated in an independent set of unrelated African ancestry (Afr) individuals from the UKB.

**Fig. S5.**
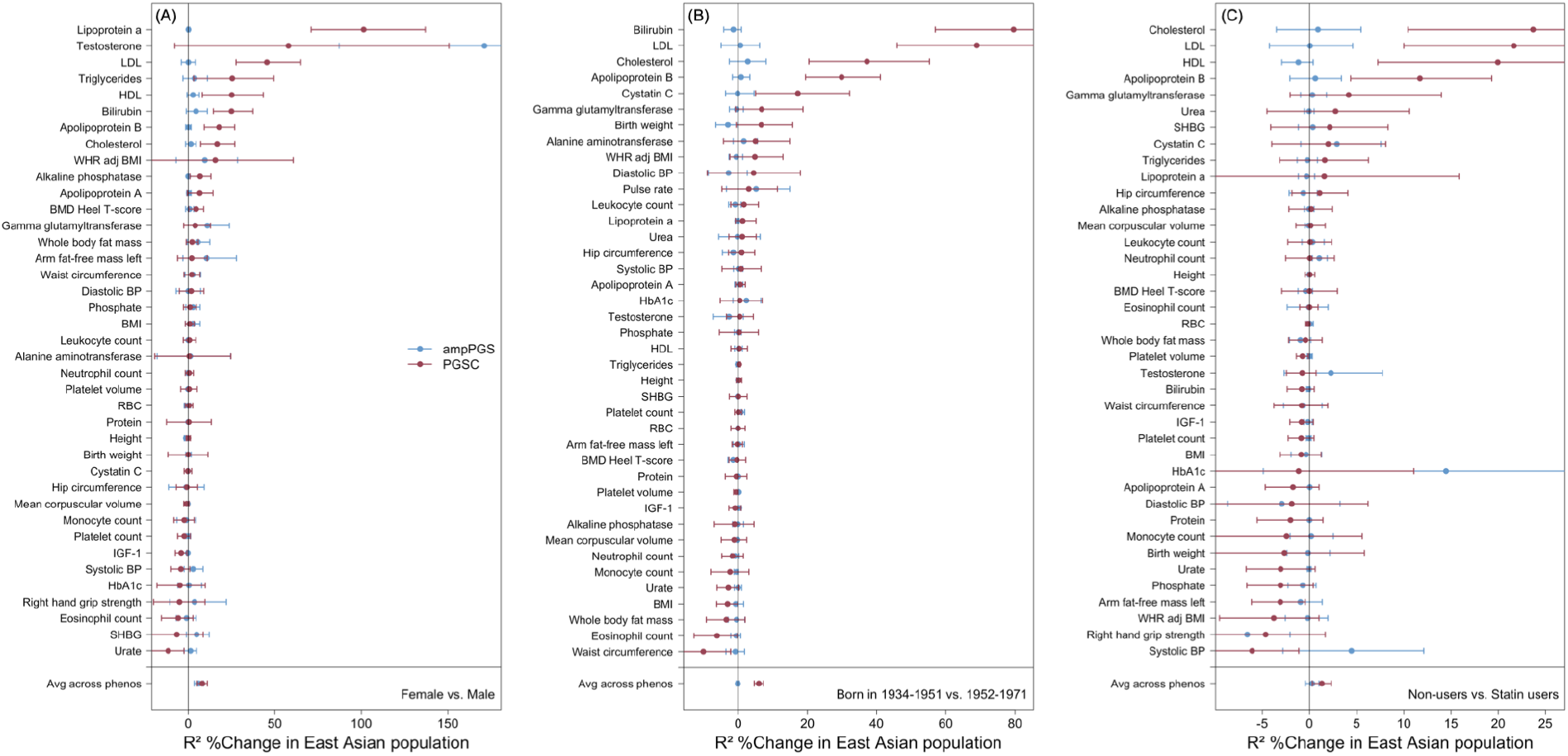
PGSC improves phenotype prediction across contexts in the UKB East Asian population. R^2^ %Change for phenotype prediction achieved by PGSC (red) and ampPGS (blue) relative to PGS across 39 quantitative traits, for contexts (A) sex, (B) age, and (C) statin treatment status. The x-axis shows the relative change in incremental R^2^, while the y-axis lists individual phenotypes, with the average improvement across all traits shown at the bottom. Error bars represent 95% confidence intervals computed from 10,000 bootstrap samples. All scores were trained on a set of unrelated WB individuals and evaluated in an independent set of unrelated East Asian ancestry (Asn) individuals from the UKB.

**Fig. S6.**
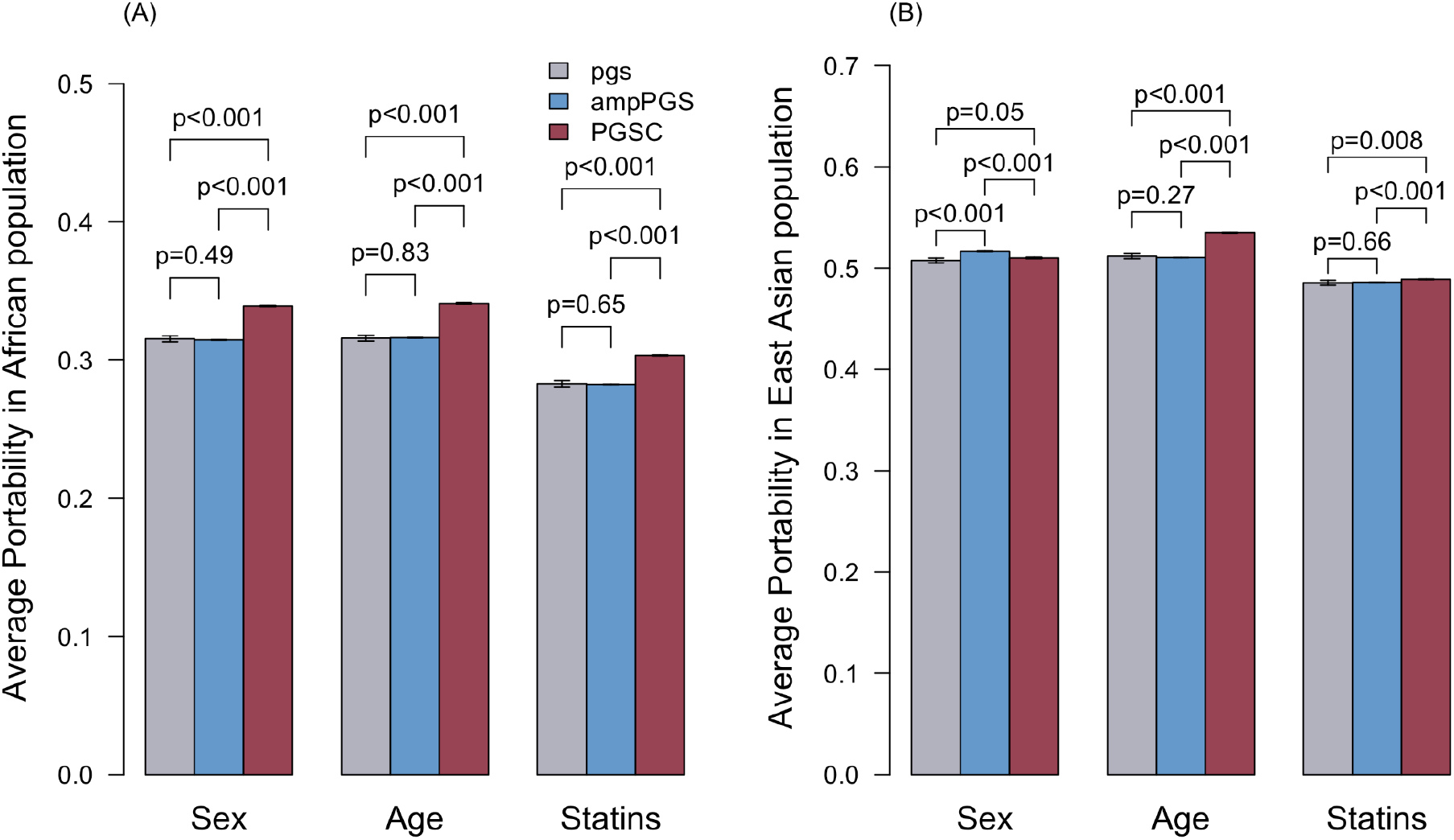
Portability comparison of PGSC to ampPGS and the standard PGS. Average score portability into the (A) Afr and (B) Asn UKB populations, for each model: PGSC (red), ampPGS (blue), and the standard PGS (grey), across three contexts: sex, age, and statin use. Bars show the mean portability averaged across all included phenotypes; error bars are 95% confidence intervals, with two-sided p-values per model comparison notated above. Portability is defined as the relative R^2^ per target population compared to the Euro population.

**Fig. S7.**
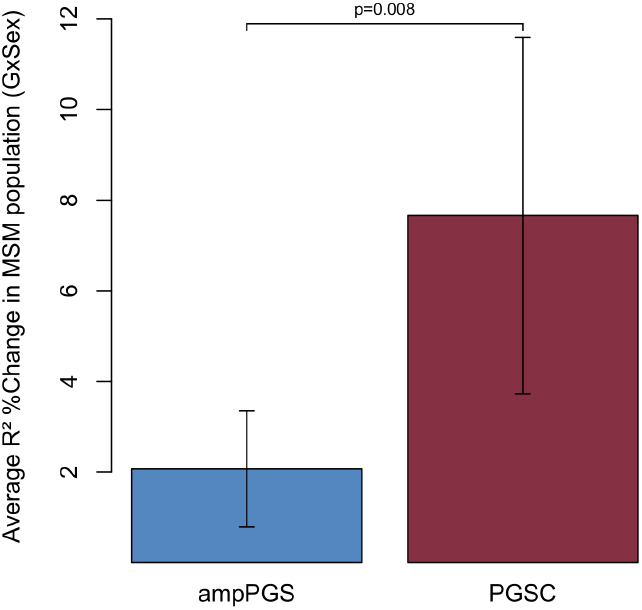
Avg R^2^ %Change in MSM given GxSex effects. Average R^2^ %Change for phenotype prediction achieved by PGSC (red) and ampPGS (blue) relative to PGS across 16 quantitative traits given GxSex effects. Error bars represent 95% confidence intervals computed from 10,000 bootstrap samples. All scores were trained on a set of unrelated WB individuals and evaluated in an independent set of 18,493 unrelated individuals in MSM genetically similar to European ancestry reference groups (as defined in Belbin 2021^21^).

**Fig. S8.**
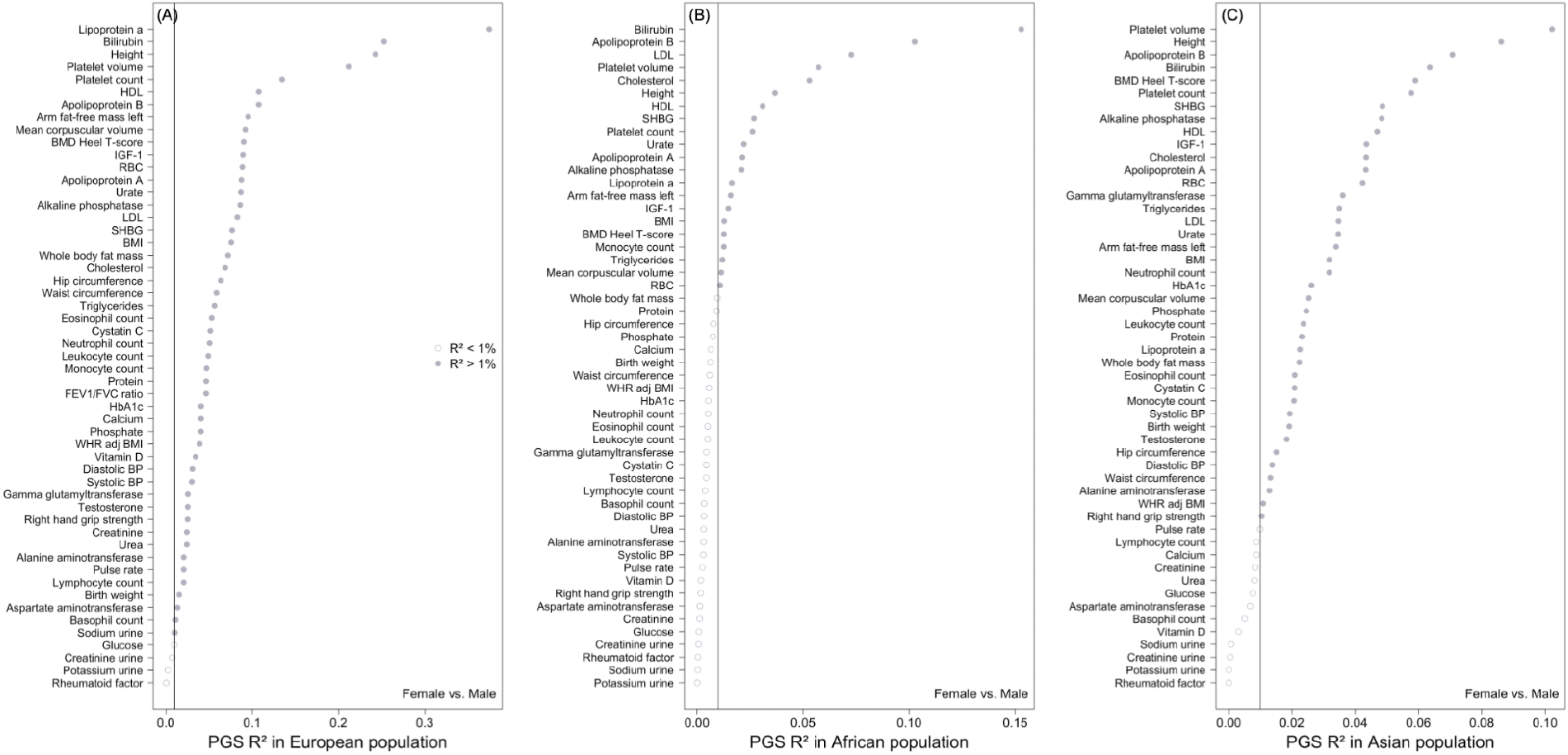
Standard PGS R^2^ in the UKB populations. PGS R^2^ across 53 quantitative traits for (A) European, (B) African, and (C) East Asian ancestry UKB populations. The x-axis shows the R^2^, while the y-axis lists individual phenotypes. All scores were trained on a set of unrelated WB individuals. The vertical black line indicates the 0.01 PGS R^2^ threshold required for phenotype inclusion in PGSC and ampPGS analyses.

**Fig. S9.**
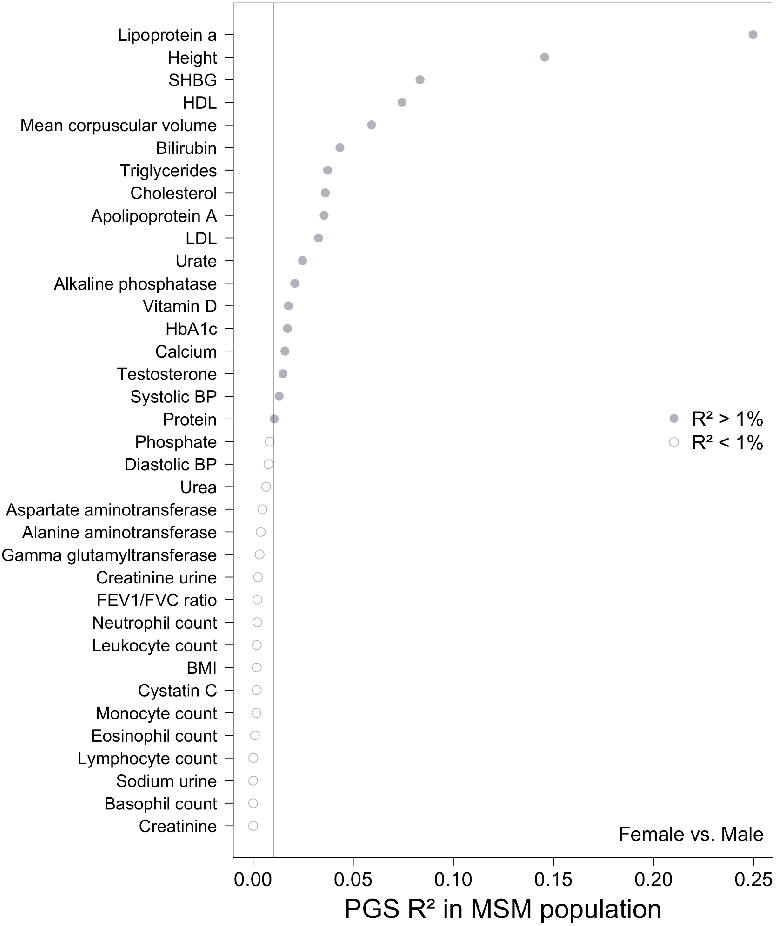
Standard PGS R^2^ in MSM population. PGS R^2^ across 36 quantitative traits in MSM. The x-axis shows the R^2^, while the y-axis lists individual phenotypes. All scores were trained on a set of unrelated WB individuals. The vertical black line indicates the 0.01 PGS R^2^ threshold required for phenotype inclusion in both PGSC and ampPGS analyses.

**Fig. S10.**
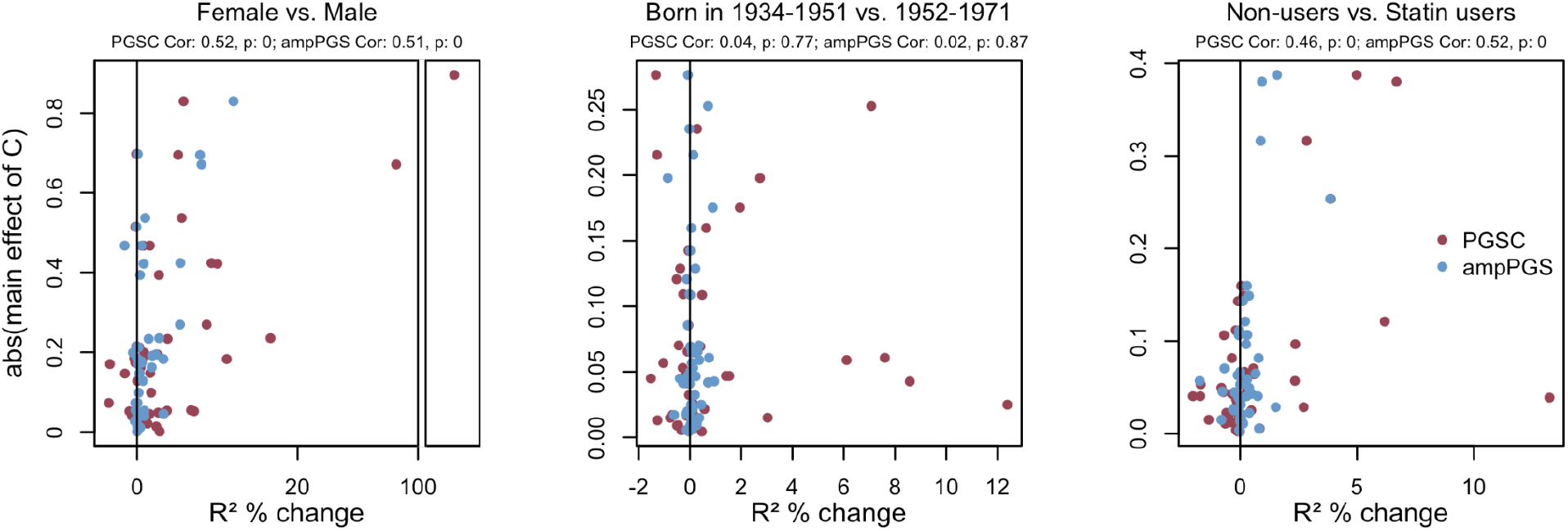
R^2^ %Change in UKB Euro population versus the main effect of C. Each panel plots the main effect of the respective context on a phenotype against the R^2^ %Change for the PGSC (red) and ampPGS (blue) in the Euro population relative to the baseline PGS, for three contexts: sex, age, and statin use. Text above each panel reports the Pearson correlation coefficient and associated two-sided p-value. The sex panel includes an x-axis break to better display testosterone without compressing the remaining points.

## Supplementary tables

**Table S1** | **Studies reporting prediction results from context-dependent polygenic scores**. “GxC model” indicates whether the model incorporates locus-specific interactions or PGS-level interactions. “GxC claim” indicates whether the study aims for improved out-of-sample prediction over an additive model or statistically significant in-sample improvement of model fit. “Scale robust” refers to testing if prediction gains are robust to a secondary phenotype scale (e.g., a log transformation). [1] The mash-based model applied to context-specific effects consistently outperforms the standard PGS, but this is confounded by benefits from Bayesian shrinkage unrelated to context-specific effects^10,23^. (In Zhu 2023, the simpler sex-specific PGS outperforms the typical PGS for 1/27 traits: testosterone.) [2] Prediction improvement is not shown to be statistically significant (confidence intervals are bootstrapped from the training data)^61^. [3] GxEprs predictions are not truly out of sample because they are re-tuned using individuals in the validation cohort^36,37^; as a result, they could not be applied, for example, to a newly genotyped individual in a clinical setting. [4] PGS-context interaction tests are commonly used to detect polygenic GxC and are reviewed in Herrera-Luis 2024^11^. [5] The demonstrated improvement in prediction accuracy refers to in-sample model fit; prediction is not tested in held-out data^26^.

**Table S2** | **List of UKB phenotypes included in analysis**.

Lists all 64 UKB quantitative phenotypes analyzed, with their field codes, including 11 log-transformed phenotypes and the constructed WHRadjBMI phenotype (based on Zhu et al. 2023^10^).

**Table S3** | **UKB population counts by context**.

Reports individual counts for each context category broken down by all four UKB populations included.

**Table S4** | **PGSC, ampPGS, and PGS R**^**2**^**s in UKB Euro population given sex as the context**.

Reports PGS R^2^ and p-value, plus ampPGS and PGSC R^2^, bootstrap R^2^ Change with 95% CIs, and best-fit p-value thresholds per phenotype. Also includes fitted model weights and each context’s standardized effect on the phenotype.

**Table S5** | **PGSC, ampPGS, and PGS R**^**2**^**s in UKB Euro population given age as the context**.

Reports PGS R^2^ and p-value, plus ampPGS and PGSC R^2^, bootstrap R^2^ Change with 95% CIs, and best-fit p-value thresholds per phenotype. Also includes fitted model weights and each context’s standardized effect on the phenotype.

**Table S6** | **PGSC, ampPGS, and PGS R**^**2**^**s in UKB Euro population given statins as the context**.

Reports PGS R^2^ and p-value, plus ampPGS and PGSC R^2^, bootstrap R^2^ Change with 95% CIs, and best-fit p-value thresholds per phenotype. Also includes fitted model weights and each context’s standardized effect on the phenotype.

**Table S7** | **List of GENIE UKB phenotypes included in analysis**.

Lists all 33 UKB quantitative phenotypes included in GxC and heteroskedasticity analyses (sourced from Pazokitoroudi et al. 2024^8^), each matched to its corresponding PGSC UKB phenotype.

**Table S8** | **PGSC, ampPGS, and PGS R**^**2**^**s in UKB Afr population given sex as the context**. Reports PGS R^2^ and p-value, plus ampPGS and PGSC R^2^, bootstrap R^2^ Change with 95% CIs, and best-fit p-value thresholds per phenotype. Also includes fitted model weights and each context’s standardized effect on the phenotype.

**Table S9** | **PGSC, ampPGS, and PGS R**^**2**^**s in UKB Afr population given age as the context**. Reports PGS R^2^ and p-value, plus ampPGS and PGSC R^2^, bootstrap R^2^ Change with 95% CIs, and best-fit p-value thresholds per phenotype. Also includes fitted model weights and each context’s standardized effect on the phenotype.

**Table S10** | **PGSC, ampPGS, and PGS R**^**2**^**s in UKB Afr population given statins as the context**.

Reports PGS R^2^ and p-value, plus ampPGS and PGSC R^2^, bootstrap R^2^ Change with 95% CIs, and best-fit p-value thresholds per phenotype. Also includes fitted model weights and each context’s standardized effect on the phenotype.

**Table S11** | **PGSC, ampPGS, and PGS R**^**2**^**s in UKB Asn population given sex as the context**.

Reports PGS R^2^ and p-value, plus ampPGS and PGSC R^2^, bootstrap R^2^ Change with 95% CIs, and best-fit p-value thresholds per phenotype. Also includes fitted model weights and each context’s standardized effect on the phenotype.

**Table S12** | **PGSC, ampPGS, and PGS R**^**2**^**s in UKB Asn population given age as the context**.

Reports PGS R^2^ and p-value, plus ampPGS and PGSC R^2^, bootstrap R^2^ Change with 95% CIs, and best-fit p-value thresholds per phenotype. Also includes fitted model weights and each context’s standardized effect on the phenotype.

**Table S13** | **PGSC, ampPGS, and PGS R**^**2**^**s in UKB Asn population given statins as the context**.

Reports PGS R^2^ and p-value, plus ampPGS and PGSC R^2^, bootstrap R^2^ Change with 95% CIs, and best-fit p-value thresholds per phenotype. Also includes fitted model weights and each context’s standardized effect on the phenotype.

**Table S14** | **Cross-population portability comparison of PGSC vs PGS**.

For each context in the Afr and Asn populations, this table reports the Pearson correlation and p-value between the per-phenotype R^2^ %Change in that population and the corresponding gain in the Euro population, computed over each set of phenotypes with PGS R^2^ > 0.01 in both the target and Euro populations.

**Table S15** | **List of MSM phenotypes included in analysis**.

Lists the 36 MSM biobank phenotypes used for replication, each matched to its corresponding UKB phenotype, along with the median and total number of observations.

