## Supplementary Information for "Locus-specific gene-context interactions improve polygenic prediction"

August 26, 2026

#### 1 Capturing Heteroskedasticity with PGSC

For a binary context, the per-context estimates under the additive model with heteroskedasticity can be written as:

$$\hat{\beta}_a \sim \mathcal{N}(\beta, \Sigma_a); \quad \hat{\beta}_b \sim \mathcal{N}(\beta, \Sigma_b)$$

where $\hat{\beta}_a$ and $\hat{\beta}_b$ are independent estimates of the genetic effect per context, and $\Sigma_a$ and $\Sigma_b$ are their respective covariance matrices. For simplicity, we assume that $\Sigma_c = \frac{s_c^2}{f_c} D$, where $D$ is a diagonal matrix inversely proportional to the per-SNP heterozygosity, $f_c$ is the fraction of individuals in context $c$, and $s_c^2$ is the residual phenotypic variance in context $c$.

The PGSC prediction with context-nonspecific GxC weight $\bar{\rho} = \rho_a = \rho_b$ is:

$$\begin{aligned} PGSC &= g\hat{\beta} + \bar{\rho}(g\hat{\lambda}) \\ &= g\left(f_a\hat{\beta}_a + f_b\hat{\beta}_b\right) + \bar{\rho} g\sqrt{f_a f_b}\left(\hat{\beta}_a - \hat{\beta}_b\right) \\ &= g\left((f_a + \sqrt{f_a f_b}\bar{\rho})\hat{\beta}_a + (f_b - \sqrt{f_a f_b}\bar{\rho})\hat{\beta}_b\right) \end{aligned}$$

where $\hat{\beta}$ and $\hat{\lambda}$ are the additive and GxC estimates from GxC GWAS (under the homoscedastic model), assuming that the context $C$ is encoded to mean 0 and variance 1.

Then the PGSC prediction MSE is:

$$\begin{aligned} \mathbb{E}\left((g\beta - PGSC(\bar{\rho}))^2\right) &= \mathbb{E}\left(\left(g\beta - \left(g\hat{\beta} + \bar{\rho}(g\hat{\lambda})\right)\right)^2\right) \\ &= \mathbb{V}\left((f_a + \sqrt{f_a f_b}\bar{\rho})\hat{\beta}_a + (f_b - \sqrt{f_a f_b}\bar{\rho})\hat{\beta}_b\right) \\ &= (f_a + \sqrt{f_a f_b}\bar{\rho})^2 \operatorname{tr}(\Sigma_a) + (f_b - \sqrt{f_a f_b}\bar{\rho})^2 \operatorname{tr}(\Sigma_b) \\ &\propto (f_a + \sqrt{f_a f_b}\bar{\rho})^2 \frac{s_a^2}{f_a} + (f_b - \sqrt{f_a f_b}\bar{\rho})^2 \frac{s_b^2}{f_b} \end{aligned}$$

where $g$ is integrated out, assuming genotypes are standardized to mean zero and variance one. Minimizing with respect to $\bar{\rho}$ gives the optimal weight on the GxC effect estimate under the additive model with heteroskedasticity:

$$\arg\min_{\bar{\rho}} \mathbb{E}\left((g\beta - PGSC(\bar{\rho}))^2\right) = \sqrt{f_a f_b}\frac{s_b^2 - s_a^2}{f_b s_a^2 + f_a s_b^2}$$

### 2 Simulations

#### 2.1 Simulating genotypes and phenotypes with GxC interaction effects

We simulate genotypes $G$ for $N$ independent individuals to assess how PGSC performs while varying heritability, GxC heritability, population size, etc. For each of $S$ SNPs, we draw a minor allele frequency (MAF) independently from a uniform distribution,

$$\text{MAF}_s \overset{\text{iid}}{\sim} \mathcal{U}(0.05,\, 0.5),$$

and simulate individual genotypes as the count of minor alleles across two independent draws,

$$G_{is} \overset{\text{ind}}{\sim} \text{Bin}(2,\, \text{MAF}_s).$$

We scale each SNP to have mean 0 and variance 1, and set the number of causal SNPs ($S_{caus}$) to 10% of $S$. Effect sizes $\beta_s$ and $\lambda_s$ are set to zero at all non-causal SNPs.

We simulate phenotypes according to the following model:

$$y = G\beta + \frac{\gamma}{\sigma_g}(G\beta) * C + (G\lambda) * C + C\alpha + (\sigma_a 1_a + \sigma_b 1_b) * \epsilon. \tag{1}$$

where

$y \in \mathbb{R}^{N\times 1}$: phenotype vector for $N$ individuals

$C \in \left\{\frac{-f_a}{\sqrt{f_a f_b}}, \frac{f_b}{\sqrt{f_a f_b}}\right\}^{N\times 1}$: binary context vector, normalized to mean 0 and variance 1; $f_a$ and $f_b$ are the fractions of individuals in each context

$G \in \mathbb{R}^{N\times S}$: scaled genotype matrix on $N$ individuals at $S$ SNPs

$\beta \in \mathbb{R}^{S\times 1}$, $\beta_s \overset{\text{iid}}{\sim} \mathcal{N}\left(0, \frac{1}{S_{caus}}\sigma_g^2\right)$ at causal SNPs: additive genetic effect ($\beta_s = 0$ for non-causal SNPs), where $\sigma_g^2$ is the additive heritability

$\gamma \in \mathbb{R}$: genome-wide amplification GxC effect; normalized by $\frac{1}{\sigma_g}$

$\lambda \in \mathbb{R}^{S\times 1}$, $\lambda_s \overset{\text{iid}}{\sim} \mathcal{N}\left(0, \frac{1}{S_{caus}}\sigma_{g\times c}^2\right)$ at causal SNPs: locus-specific GxC effect ($\lambda_s = 0$ for non-causal SNPs), where $\sigma_{g\times c}^2$ is the locus-specific GxC heritability

$\alpha$: context main effect on $y$

$1_a, 1_b \in \{0,1\}^{N\times 1}$: binary context indicators, with $1_{b,i} = 1$ if $i \in b$ and $1_{a,i} = 1 - 1_{b,i}$

$\sigma_a = \sqrt{\frac{2\sigma_\epsilon^2}{1+\eta}}$; $\sigma_b = \sigma_a\sqrt{\eta}$ where $\eta$ is the heteroskedasticity parameter ($\eta \in \mathbb{R}_{>0}$), and $\sigma_\epsilon^2$ is phenotypic variance unaccounted for by genetic or GxC heritability ($\sigma_\epsilon^2 = 1 - \sigma_g^2 - \gamma^2 - \sigma_{g\times c}^2$); when $\eta = 1$, $\sigma_a = \sigma_b = \sigma_\epsilon$ (homoskedasticity)

$\epsilon \sim \mathcal{N}(0, I_N)$: noise vector

#### 2.2 Phenotype scale transformation

To assess robustness to phenotype scale, we apply a Box-Cox power transformation to phenotypes simulated from (1). Prior to transformation we shift $y$ to ensure it's $\geq 1$,

$$y' = y - \min(y) + 1,$$

and then apply the Box-Cox transform with coefficient $\lambda_{\text{Box-Cox}}$:

$$y'' = \begin{cases} \dfrac{y'^{\lambda_{\text{Box-Cox}}} - 1}{\lambda_{\text{Box-Cox}}} & \lambda_{\text{Box-Cox}} \neq 0, \\ \log(y') & \lambda_{\text{Box-Cox}} = 0. \end{cases}$$

The transformed phenotype is then standardized to mean 0 and variance 1. When $\lambda_{\text{Box-Cox}} = 1$ (the default), the transformation is equivalent to the original simulation. This analysis follows the framework in Costantino et al. 2026.

#### 3 PGSC Algorithm

**Algorithm 1:** PGSC Construction

**Input :** y = phenotypes, G = genotypes, X = covariates, C = context

**Output :** PGSC

**Training: Perform a GWAS**

$$y \sim X\alpha + C\gamma + G_j\beta_j + \epsilon$$

**Training: Perform a context-specific GWAS**

$$y \sim X\alpha + C\gamma + G_j\beta_j + (G_j * C)\lambda_j + \epsilon$$

$$\triangleright\ C_i = \begin{cases} C_a & \texttt{if } i \in a \\ C_b = -\dfrac{C_a n_a}{n_b} & \texttt{if } i \in b \end{cases}$$

**Testing: Optimize PGS parameter** ($\tau_g$)

$$PGS = G\hat{\beta}_{\tau_g}$$

▷ $\tau_g, \tau_{g\times c}$: `p-value thresholds`

**Testing: Optimize PGSC parameters** ($\tau_{g\times c}, \rho_a, \rho_b$)

$$PGSC = PGS + (G\hat{\lambda}_{\tau_{g\times c}}) * (\rho_a 1_a + \rho_b 1_b)$$

▷ $1_a, 1_b$: `context-group indicator vectors`

**Validation: Build a PGS**

$$PGS = G\hat{\beta}_{\tau_g}, \quad R^2(y, PGS)$$

▷ `Implementing optimized` $\tau_g$

**Validation: Build a PGSC**

$$PGSC = PGS + (G\hat{\lambda}_{\tau_{g\times c}}) * (\rho_a 1_a + \rho_b 1_b)$$

▷ `Implementing optimized` $\tau_g$, $\tau_{g\times c}$, $\rho_a$, $\rho_b$

$$R^2(y, PGSC)$$

This algorithm is valid for any choice of $C_a$, since $C$ will always have mean zero. Our choice of $C_a = \frac{f_b}{\sqrt{f_a f_b}}$ additionally ensures that $C$ has variance 1, enabling the simple formula in Main Eq 1 for the optimal $\bar{\rho}$ for the additive model with heteroskedasticity.
